# High-throughput functional genomics and transcriptomics reveal molecular mechanisms regulating the synergistic fungicidal activity of copper and zinc

**DOI:** 10.64898/2026.10.02.756351

**Authors:** Joshua D. Kerkaert, Renato Carvalho, Ruby E. Schnirman, Christine Xin Yan Wu, Brandon Reyes-Chavez, Yuxin Zhang, Margaret K. A. Koker, Louisa M. Smieska, Lori B. Huberman

## Abstract

Fungal diseases cause devastating crop infestations and millions of deaths annually. To improve disease treatment development, we must understand the biological processes these treatments target and the fungal cellular responses they elicit. Current methods to identify genetic responses of fungi to antifungal drugs and fungicides typically involve laborious molecular genetics experiments or transcriptomics. We utilized high-throughput screens in the basidiomycete yeast *Rhodotorula* (*Rhodosporidium*) *toruloides* to rapidly identify genes associated with the fungal response to a copper-zinc fungicide developed by VM Agritech and compared identified genes to those transcriptionally activated in response to the fungicide in *R. toruloides* and the filamentous ascomycete *Neurospora crassa*. These ‘omics technologies combined with molecular genetics and biochemical analyses showed copper and zinc synergize to inhibit fungal growth due, at least in part, to an increase in cellular zinc concentrations when copper is present. Intriguingly, although we identified genes involved in the copper-zinc fungicide response using both high-throughput screens and transcriptomics, there was little overlap in genes identified through the two methods. Our findings demonstrate the power of combining functional genomics and transcriptomics approaches in identifying a more comprehensive map of the genetic mechanisms necessary for responding to fungicides.

## INTRODUCTION

Fungal diseases of plants and humans result in estimated crop losses of 20-30% and millions of deaths annually^1,2^. Unfortunately, available clinical antifungals and agricultural fungicides are limited, and their continued use is threatened by antifungal resistance^3^. Compounding this problem, several agricultural fungicides and clinical antifungals have the same mechanism of action, resulting in environmentally acquired cross-resistance^3–5^. Thus, we must develop new disease management strategies, fungicides, and antifungal drugs.

Since their development in 1885, copper-based fungicides have been one of the widest used chemical classes for controlling fungal, bacterial, and viral pathogens in conventional and organic farming^3,6,7^. Toxic copper concentrations lead to reactive oxygen species generation and protein mismetallation, misfolding, and aggregation^8–14^. This multimodal mechanism of action decreases resistance development relative to single target fungicides and is a significant appeal of copper-based fungicides. However, soil copper accumulation following long-term copper fungicide use is detrimental to plant health^15^.

Like copper, zinc is an essential micronutrient with antimicrobial properties at high concentrations. Zinc also has a multimodal mechanism of toxicity through protein mismetallation, generation of reactive oxygen species, and altering transport of other metal micronutrients^16–21^. While zinc oxide nanoparticles can protect against fungal and bacterial pathogens^22,23^, zinc is not widely used as a fungicide. Rather, application of zinc sulfate and zinc chelates is used to correct zinc deficiencies, a common problem in soils^24,25^.

The company VM Agritech developed a novel copper fungicide-zinc fertilizer formulation. Given that the metals have overlapping mechanisms of antifungal activity and zinc is known to alter the transport of other metals, we hypothesized the combination of copper and zinc may synergize or antagonize fungicide efficacy^8,9,16–18^. We explored interactions between copper and zinc using two model fungal systems, the ascomycete filamentous fungus *Neurospora crassa* and the basidiomycete yeast *Rhodotorula* (*Rhodosporidium*) *toruloides*.

Our data show copper and zinc interact synergistically to inhibit fungal growth, and this synergy is at least partially accomplished through increased cellular zinc concentrations when copper is present. Characterization of the transcriptional responses to copper and zinc alone and in combination revealed that only a small number of genes exhibited synergy at the transcriptional level. To identify genes required for resistance to copper and zinc alone and in combination, we used a combination of transcriptomics and high-throughput screens of an *R. toruloides* barcoded insertional mutagenesis library^26^. We identified mechanisms of metal resistance using both methods, but there was very limited overlap, suggesting that transcriptomics and functional genomics are complementary in determining the genetic mechanisms of stress resistance in fungi.

## RESULTS

### Copper and zinc synergize to inhibit fungal growth

VM Agritech developed Curezin^®^, a mixture of copper sulfate and zinc sulfate at a ratio of 1 Cu^2+^:0.96 Zn^2+^ in an ammonium-phosphate buffer, as a broad-spectrum fungicide that doubles as a zinc fertilizer. Because copper and zinc toxicity have similar modes of action^8–14,16–21^, we hypothesized there may be interactions in the fungal response to these two metals. To test this hypothesis, we first quantified growth of two fungal species: a model Ascomycete filamentous fungus, *Neurospora crassa*, and a Basdiomycete yeast, *Rhodotorula (Rhodosporidium) toruloides* in the presence of copper sulfate and zinc sulfate in an ammonium-phosphate buffer individually and in combination. These species represent two major fungal phyla containing many critical plant, human, and animal pathogens, as well as two of the major fungal morphological lifestyles.

Treatment with copper alone caused almost complete growth inhibition in both species at 10 mM relative to cells grown in media alone or media containing only buffer (Fig 1A-1B). *N. crassa* growth was almost completely inhibited at 9.56 mM zinc (the highest soluble concentration of ZnSO_4_ in Vogel’s minimal medium^27^), while *R. toruloides* growth was only reduced by 10% (Fig 1A, 1B, and S1A-S1C). Treatment of *N. crassa* and *R. toruloides* with copper and zinc in combination resulted in a greater growth reduction than the sum effect of treatment with either compound individually (Fig 1 and S1). In *N. crassa*, treatment with 1.91 mM zinc resulted in 4.4% growth inhibition, and treatment with 2 mM copper caused 18.9% growth inhibition. However, a mixture of 1.91 mM zinc with 2 mM copper caused 77.3% growth inhibition (Fig 1A and S1A). Similarly, treatment of *R. toruloides* with 9.56 mM zinc and 3.33 mM copper caused 10.1% and 13.4% growth inhibition, respectively, while treatment with the combination of the two resulted in 70.5% growth inhibition (Fig 1B and S1B-S1C). To determine if the copper-zinc interaction was additive or synergistic, we utilized four interaction models available in the SynergyFinder Plus^28^ tool: zero-interaction potency^29^, highest single agent, Bliss independence, and Loewe additivity. All four models classified the copper-zinc interaction as synergistic in both *N. crassa* and *R. toruloides* (Fig 1C, 1D, and S1D-S1I).

**Fig 1.**
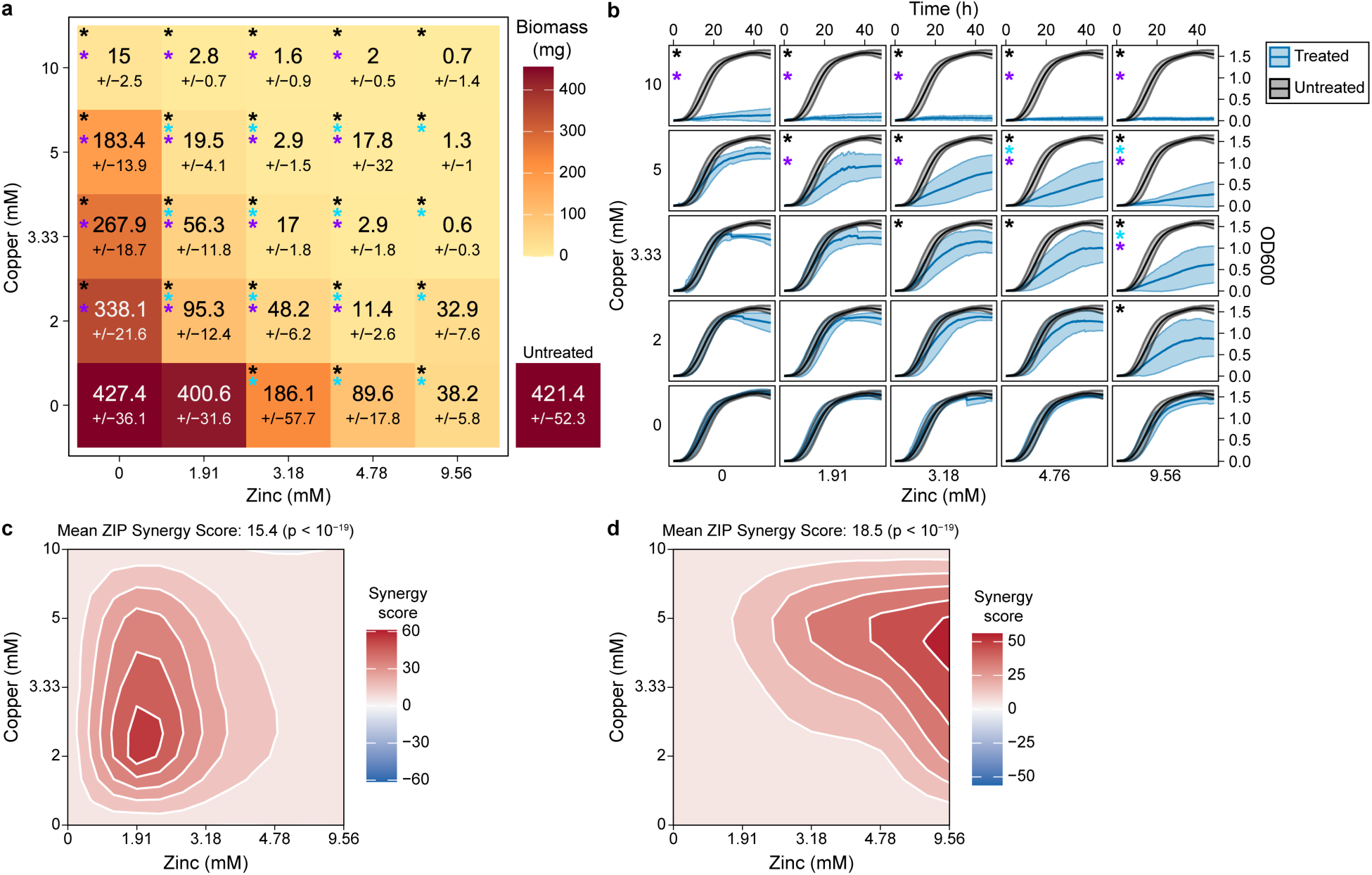
Copper and zinc act synergistically to inhibit fungal growth. **A)** Biomass accumulation 2 d after direct inoculation of wild type *N. crassa* into 100 ml of minimal medium containing the indicated concentrations of copper sulfate and zinc sulfate in an ammonium-phosphate buffer. The top numbers in each square are the mean and the lower numbers are the standard deviation of four biological replicates. **B)** Growth curves (OD600) of wild type *R. toruloides* grown in media alone (black lines) or media with the indicated concentrations of copper sulfate and zinc sulfate in an ammonium-phosphate buffer (blue lines). Lines are averages and colored bands are standard deviations of six biological replicates. **C-D)** Zero-interaction potency (ZIP) model^29^ synergy plot for the percent inhibition of (**C**) *N. crassa* and (**D**) *R. toruloides* by the indicated concentration of copper sulfate and zinc sulfate. To generate percent inhibition values, the (**C**) *N. crassa* biomass data (from **A**, shown in Fig S1A) and (**D**) *R. toruloides* area under the curve values (from **B**, shown in Fig S1B-C) were normalized to the untreated control for each experiment. The normalized data were used for synergy calculations and contour plot generation using the SynergyFinder+ R package^28^. *p < 0.05 as determined by either a one-way ANOVA with a Dunnett’s post-hoc test with untreated as the control group (black stars) or a two-way ANOVA with a Tukey’s post-hoc test on biomass (**A**) or area under the curve (**B**, shown in Fig S1B) (blue and purple stars). Colored stars indicate statistically significant differences in growth in the respective treatment relative to: untreated cells (black), treatment with the same concentration of copper alone (blue), or treatment with the same concentration of zinc alone (purple).

Zinc is known to alter the transport of other metals^17,18^. Thus, we hypothesized the synergistic activity of copper and zinc was due to increased cellular copper when the metals were applied in combination. To test this hypothesis, we quantified relative cellular metal content using x-ray fluorescence imaging. *R. toruloides* cells were incubated in minimal media supplemented with ammonium-phosphate buffer, 5 mM copper, 4.78 mM zinc, or 5 mM copper and 4.78 mM zinc and then washed with minimal media to remove free metals. Surprisingly, the cellular copper concentration was 40% lower in dual-treated cells compared to copper alone (Fig 2A and S2). In contrast, cellular zinc concentration increased by 74% in dual-treated cells compared to cells treated with zinc alone (Fig 2B and S2). These data suggest the synergy of copper and zinc treatment may be due to increased cellular zinc.

**Fig 2.**
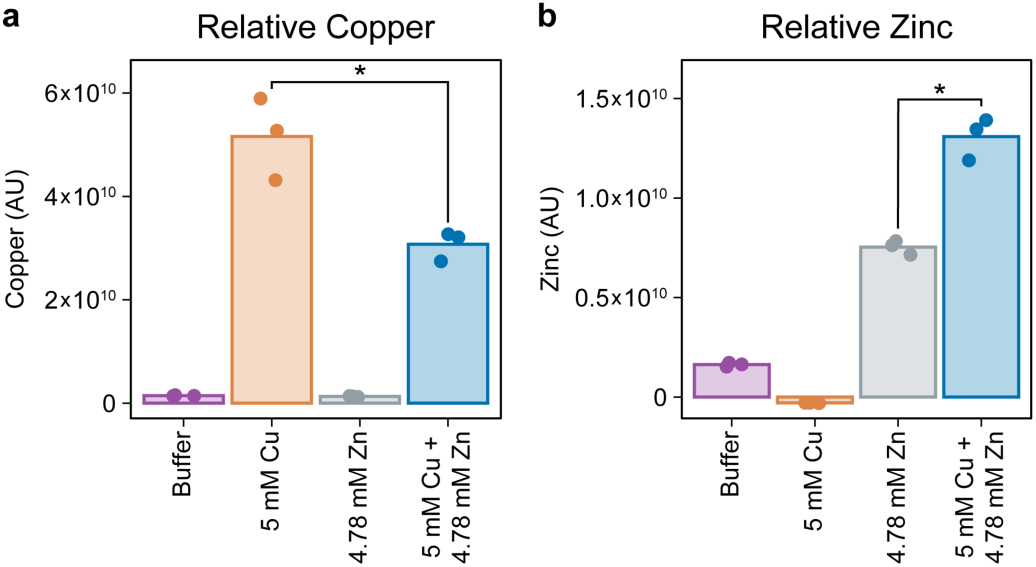
Treatment of *R. toruloides* with copper and zinc in combination increases cellular zinc concentrations and decreases cellular copper concentrations compared to zinc or copper treatment alone, respectively. Relative cellular concentrations of **(A)** copper and **(B)** zinc quantified using x-ray fluorescence imaging on an equal amount of *R. toruloides* cells grown in media supplemented with ammonium-phosphate buffer or ammonium-phosphate buffer with 5 mM copper, 4.78 mM zinc, or 5 mM copper and 4.78 mM zinc. Bars are the averages and dots are the individual data points of three biological replicates. *p < 0.01 by one-way ANOVA with a Tukey’s post-hoc test. AU = arbitrary units.

High concentrations of copper and zinc individually are fungicidal^30–33^. However, it is possible combining the two metals altered fungicidal activity. To address this possibility, we performed a time-kill assay on *R. toruloides* treated with 10 mM or 20 mM copper with or without 9.56 mM zinc. Treatment with 10 mM copper resulted in a 1.4-fold decrease in recoverable colony forming units (CFU) per mL over 48 h, whereas treatment with 10 mM copper and 9.56 mM zinc yielded a 16-fold decrease in recoverable CFU/mL compared to the starting population (Fig S3). Treatment with 20 mM copper reduced the number of viable cells to below the limit of detection (50 CFU/mL) by 24 h. While 20 mM copper plus 9.56 mM zinc displayed clear fungicidal activity, with a 3,840-fold reduction in viability, we still recovered 567 CFU/mL after 24 h and 125 CFU/mL after 48 h (Fig S3). These data suggest the addition of zinc to copper slows the rate of fungal death relative to copper alone.

### N. crassa genes involved in ribosome biogenesis and metal transport exhibit a synergistic transcriptional response to copper and zinc in combination

We hypothesized the physiological synergy of copper and zinc treatment could be due to synergy in the transcriptional response to these metals. Transcriptional synergy could manifest as a transcriptional signature distinct from treatment with copper or zinc individually or synergistic changes in gene expression magnitude. To test these possibilities, we utilized 3’ RNA sequencing (RNAseq).

We performed RNAseq on *N. crassa* mycelia after a transfer to fresh media or fresh media supplemented with buffer alone, moderately inhibitory treatments (3.33 mM copper sulfate, 3.19 mM zinc sulfate, or 2 mM copper sulfate and 1.91 mM zinc sulfate), or highly inhibitory treatments (5 mM copper sulfate, 4.78 mM zinc sulfate, or 3.33 mM copper sulfate and 3.19 mM zinc sulfate) for eight hours (Fig 1A and 3A and Datasets S1-S2). No genes were at least four-fold differentially expressed between the untreated and buffer treated conditions (Dataset S2). Thus, we performed differential expression analyses relative to buffer-treated cells.

**Fig 3.**
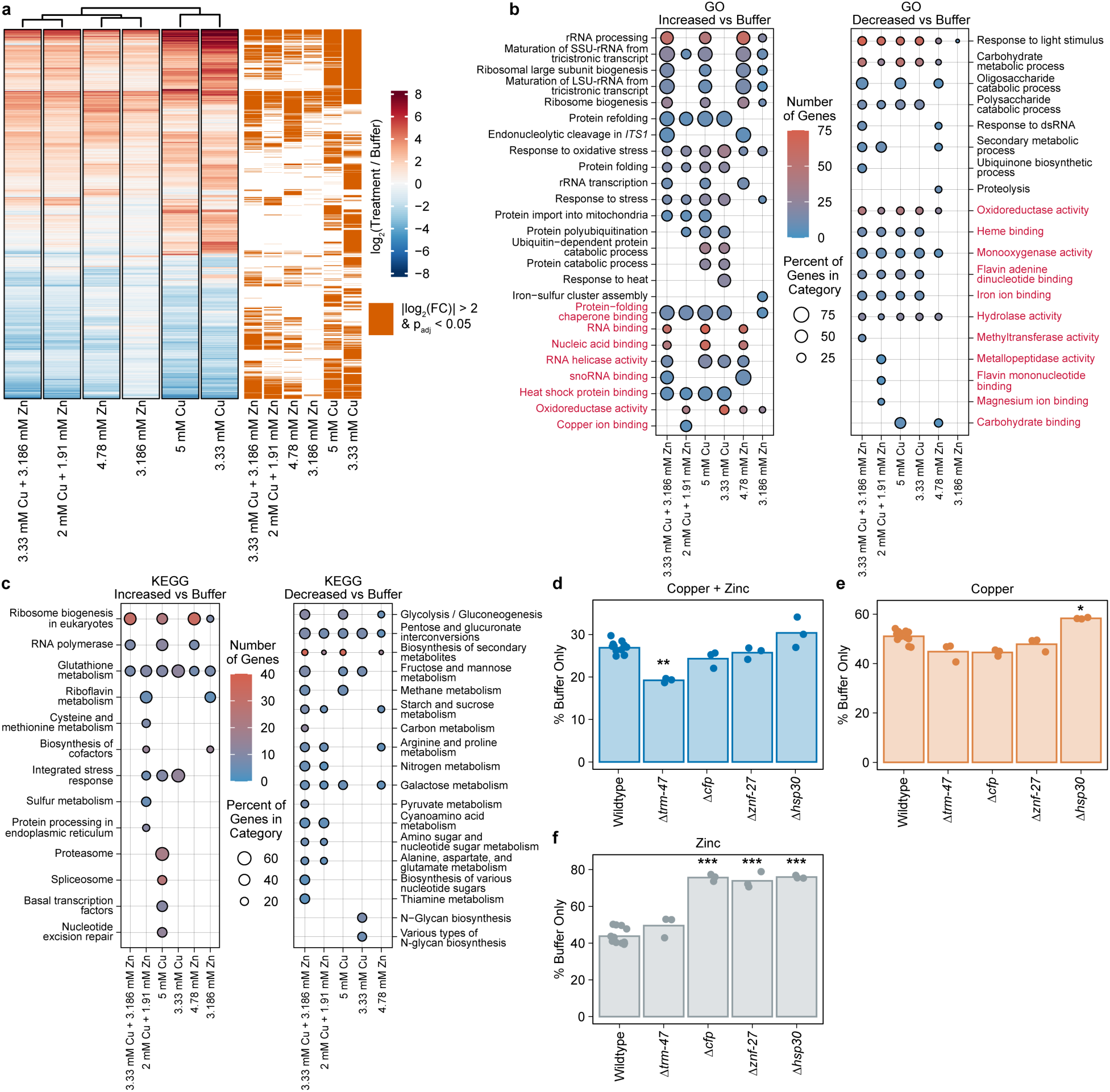
*N. crassa* upregulates protein synthesis machinery and broadly downregulates metabolism in response to copper and zinc. **A)** Heatmap of the log_2_(fold change) in gene expression during exposure to the indicated concentrations of copper sulfate and/or zinc sulfate relative to exposure with ammonium-phosphate buffer. Only genes at least four-fold differentially expressed in at least one condition relative to buffer are displayed. Orange bars to the right of the heatmap indicate genes that are at least four-fold differentially expressed in the indicated condition relative to buffer with p_adj_ < 0.05. RNAseq was performed with three biological replicates. **B-C)** (**B**) GO (Gene ontology)^34,35^ and (**C**) KEGG (Kyoto Encyclopedia of Genes and Genomes)^36–38^ functional category enrichment analyses of genes at least four-fold upregulated (left) or downregulated (right) in the indicated condition relative to buffer. Displayed comparisons had at least one significantly enriched category (p_adj_ < 0.05). Dot color represents the number of genes in the category that were differentially expressed. Dot size represents the percent of genes in the category that were differentially expressed. Text of GO biological process categories is black. Text of GO molecular function categories is red. **D-F)** Relative biomass accumulation of the indicated strains 2 d post-inoculation into media supplemented with ammonium-phosphate buffer with (**D**) 2 mM copper sulfate plus 1.91 mM zinc sulfate, (**E**) 3.33 mM copper sulfate, or (**F**) 3.19 mM zinc sulfate normalized to the biomass of the indicated strain 2 d post-inoculation into media supplemented with ammonium-phosphate buffer. Raw biomass accumulation data is in Fig S6. *p < 0.05, **p < 10^-5^, and ***p < 10^-12^ as determined by one-way ANOVA with a Dunnett’s post-hoc test compared to wild type cells. Multiple hypothesis corrections in **D-F** were performed using the growth of all strains assessed in Fig S6. Bars are the averages and dots are the data points of at least three biological replicates.

All metal treatment conditions resulted in substantial transcriptional responses relative to buffer-treated cells. Treatment with 3.33 mM copper had the largest response with 17% of *N. crassa* genes at least four-fold differentially expressed (Fig 3A and Dataset S2). Because the cellular impacts of copper and zinc (i.e. mismetallation, oxidative stress, etc.) are relatively similar, we first asked whether the transcriptional response to copper and zinc had overlapping gene sets. Only 32% of the genes differentially expressed in response to 5 mM copper were also differentially expressed in response to 4.78 mM zinc (|log_2_(fold change)| > 2, p_adj_ < 0.05) (Fig 3A and Dataset S2).

To examine the possibility that treatment with copper and zinc together regulated a unique set of genes, we compared the genes differentially expressed in the individual and dual treatments relative to buffer. Hierarchical clustering and principal component analyses suggested that while the transcriptional profile of copper plus zinc treated cells had similarity to both individual treatments, it was more similar to treatment with zinc than copper (Fig 3A and S4). In cells exposed to 3.33 mM copper and 3.19 mM zinc, 68% of genes increased and 50% of genes decreased at least four-fold in expression were also at least four-fold increased or decreased, respectively, in expression in cells treated with 3.19 mM zinc or 3.33 mM copper (Fig 3A). We also compared gene expression in cells exposed to conditions that caused similar levels of growth inhibition. In cells exposed to 2 mM copper and 1.91 mM zinc, 83% of genes increased and 76% of genes decreased at least four-fold in expression were also at least four-fold increased or decreased, respectively, in expression in cells treated with 3.19 mM zinc or 3.33 mM copper. Similarly, 72% of genes increased and 85% of genes decreased at least four-fold in expression in cells exposed to 3.33 mM copper and 3.19 mM zinc compared to buffer were also increased or decreased four-fold, respectively, in cells treated with 5 mM copper or 4.78 mM zinc (Fig 3A). Thus, the transcriptional profile of the dual-treatment conditions rejected a model of transcriptional synergy in which treatment with both copper and zinc results in a unique transcriptional signature.

We next asked if genes differentially expressed on copper and zinc individually exhibited synergy in the magnitude of the transcriptional response during exposure to the combination of copper and zinc. We compared the summed fold changes of genes increased or decreased in both treatment with 3.33 mM copper and treatment with 3.19 mM zinc individually (p_adj_ < 0.05 in both conditions and greater than four-fold changed in at least one condition) to the observed fold changes in 3.33 mM copper plus 3.19 mM zinc (Fig S4B-S4C). Genes were considered to have a synergistic transcriptional response if they were differentially expressed between the individual and dual treatments (p_adj_ < 0.05) and the magnitude of fold change in expression on copper plus zinc relative to buffer was at least 25% greater than the sum of fold changes on copper and zinc individually relative to buffer. We defined an antagonistic transcriptional response to be genes that were differentially expressed between the individual and dual treatments (p_adj_ < 0.05) where the magnitude of fold change on copper plus zinc relative to buffer was less than half the magnitude of fold change in both individual treatments relative to buffer. All other genes were considered to have no clear interaction for the purposes of this analysis.

Of 725 genes significantly increased in expression on both copper and zinc individual treatments, only 25 demonstrated synergy and 44 were antagonistic (Fig S4B). Similarly, of 260 genes whose expression was significantly decreased on both copper and zinc individually, the responses of 29 genes were synergistic and nine were antagonistic (Fig S4C). The 25 genes with synergistically increased expression included a copper-resistance associated P-type ATPase (*trm-47* [NCU04076]) and 12 genes involved in rRNA and ribosome biogenesis (Dataset S1-S2). The 44 upregulated genes with antagonistic responses to copper and zinc included *gcl-1* (NCU01157) and *cys-24* (NCU04596), encoding the catalytic and regulatory subunits, respectively, of the rate limiting enzyme in glutathione biosynthesis, and the flavohemoglobin *fhb-1* (NCU02096) involved in nitrosative stress mitigation (Dataset S1-S2). Thus, although most genes did not exhibit transcriptional synergy, a number of genes involved in protein synthesis, metal transport, and glutathione biosynthesis had either synergistic or antagonistic transcriptional responses to copper and zinc.

To test whether genes with synergistic or antagonistic transcriptional responses play roles in the physiological synergistic response to copper and zinc, we examined the resistance of deletion mutants of two upregulated genes that exhibited transcriptional synergy (*trm-47* and *aap-7* [NCU04942]) and one that exhibited transcriptional antagonism (NCU01138) to 3.33 mM copper, 3.19 mM zinc, or 2 mM copper and 1.91 mM zinc. Deletion of *aap-7* and NCU01138 had no effect on resistance to copper and zinc alone or in combination. The growth of Δ*trm-47* cells was indistinguishable from wild type cells on copper and zinc individually, but deletion of *trm-47* resulted in 30% lower biomass accumulation than wildtype in 2 mM copper plus 1.91 mM zinc, suggesting the synergistic transcriptional response of *trm-47* may play a role in the synergistic physiological response to copper and zinc in combination (Fig 3D-3F, S5, and S6).

### N. crassa upregulates genes involved in proteotoxic and oxidative stress responses and downregulates genes involved in metabolism in response to copper and zinc

Genes upregulated on copper, zinc, and the combination of copper and zinc were enriched for Gene Ontology (GO)^34,35^ and Kyoto Encyclopedia of Genes and Genomes (KEGG)^36–38^ categories associated with rRNA processing, ribosome biogenesis, and stress responses, consistent with protection from reactive oxygen species, and protein misfolding/mismetallation. Cysteine can chelate copper and zinc, and is a precursor to glutathione, which can complex with copper to stabilize its redox state and inhibit reactive oxygen species production^8,13,14^. We saw enrichment for GO and KEGG categories associated with glutathione metabolism and an upregulation of key cysteine and glutathione biosynthesis genes across all treatments (Fig 3B-3C). These genes included three glutathione S-transferases (*gst-1* [NCU05780], *gst-2* [NCU04109], and *gst-3* [NCU01320]), a cystathionine γ-lyase (*cys-16* [NCU09230]), glutamate-cysteine ligase (*gcl-1* and *cys-24*), and glutathione reductase (*gtr-1* [NCU03339])^39^.

Responses to protein denaturing stresses, such as those caused by copper, require heat shock proteins^40^ and cause increased protein turnover. Correspondingly, 12 to 14 of the 19 annotated heat shock protein encoding genes were upregulated across the copper and dual-treated conditions and five to six heat shock protein encoding genes were upregulated across the two zinc conditions. Additionally, the expression of at least five genes encoding RNA polymerase I and III subunits, responsible for rRNA, tRNA, and other small RNA transcription, at least one gene involved in every step of ribosome biogenesis, and at least 10 genes encoding 90S pre-ribosome components was upregulated in all conditions except 3.19 mM zinc, suggesting an increase in protein turnover. Genes downregulated on copper, zinc, and the combination of copper and zinc were enriched for GO and KEGG categories associated with carbohydrate and central carbon metabolism (Fig 3B-3C). However, these genes were not clustered in a particular metabolic pathway, potentially suggesting a general reduction in metabolic activity.

We hypothesized genes upregulated on copper, zinc, and the combination of copper and zinc may mediate metal resistance. To test this hypothesis, we measured biomass accumulation in 3 mL of media containing 3.33 mM copper, 3.19 mM zinc, or 2 mM copper and 1.91 mM zinc of deletion mutants of 18 additional genes with increased expression across all conditions, three additional genes that were downregulated only on copper plus zinc, and three genes that were upregulated only on copper plus zinc. From this initial screen we identified seven additional genes with potential nonredundant roles in copper and zinc resistance, which we confirmed in 100 mL cultures (Fig S5).

All seven strains grew like wild type in minimal media with buffer, except Δ*cfp* (NCU02193), encoding a pyruvate decarboxylase, and Δ*znf-27* (NCU04022), encoding a C2H2 zinc-finger transcription factor, which accumulated 17% and 15% greater biomass than wild type cells, respectively (Fig S6D). The expression of *cfp* was 4.7-fold decreased in copper plus zinc, and Δ*cfp* was 73% more resistant to 3.19 mM zinc than wild type cells (Fig 3F). Despite a greater than 6.5-fold increase in expression of *znf-27* and the heat shock protein gene *hsp30* across all conditions relative to buffer, cells lacking *znf-27* or *hsp30* were over 70% more resistant to zinc than wild type cells (Fig 3F). Critically for using copper plus zinc for disease prevention, Δ*cfp*, Δ*znf-27*, and Δ*hsp30* cells were not resistant to copper and zinc in combination (Fig 3D).

### R. toruloides increases expression of genes involved in anaplerosis and cysteine/glutathione biosynthesis in response to copper and zinc in combination

Copper and zinc also caused a synergistic growth inhibition of *R. toruloides*. To test whether the transcriptional response to copper and zinc was synergistic in *R. toruloides*, we exposed cells to minimal media or minimal media supplemented with buffer alone, 5 mM copper plus 4.78 mM zinc, 3.33 mM copper plus 3.19 mM zinc, 6.29 mM copper, 3.33 mM copper, or 9 mM zinc for 8 h prior to harvesting cells for 3’ RNAseq. No significant differences in gene expression were detected between the untreated and buffer treated conditions. Thus, all downstream analyses were performed relative to the buffer treated condition.

The transcriptional response of *R. toruloides* to copper and zinc was much less pronounced than that of *N. crassa*, so we used a two-fold change threshold for genes of interest in our analyses (Fig 4A and Datasets S3-34). The numbers of differentially expressed genes corresponded to the magnitude of the physiological response to treatment conditions. Fewer than 100 genes were differentially expressed by at least two-fold during treatment with 3.33 mM copper or 9 mM zinc, while 233, 533, and 1,416 genes were differentially expressed by at least two-fold during treatment with 3.33 mM copper plus 3.19 mM zinc, 6.29 mM copper, and 5 mM copper plus 4.78 mM zinc, respectively (Fig 4A and Dataset S4). Hierarchical clustering and principal component analyses suggested copper is the primary driver of transcriptional changes for *R. toruloides* exposed to copper plus zinc and the transcriptional profile of 5 mM copper plus 4.78 mM zinc differs from treatment with copper and zinc individually (Fig 4A and S7). However, the resistance of *R. toruloides* to zinc treatment and the small resulting transcriptional changes made interpretation of transcriptional synergy in either transcriptional profile or magnitudes of change difficult to assess.

**Fig 4.**
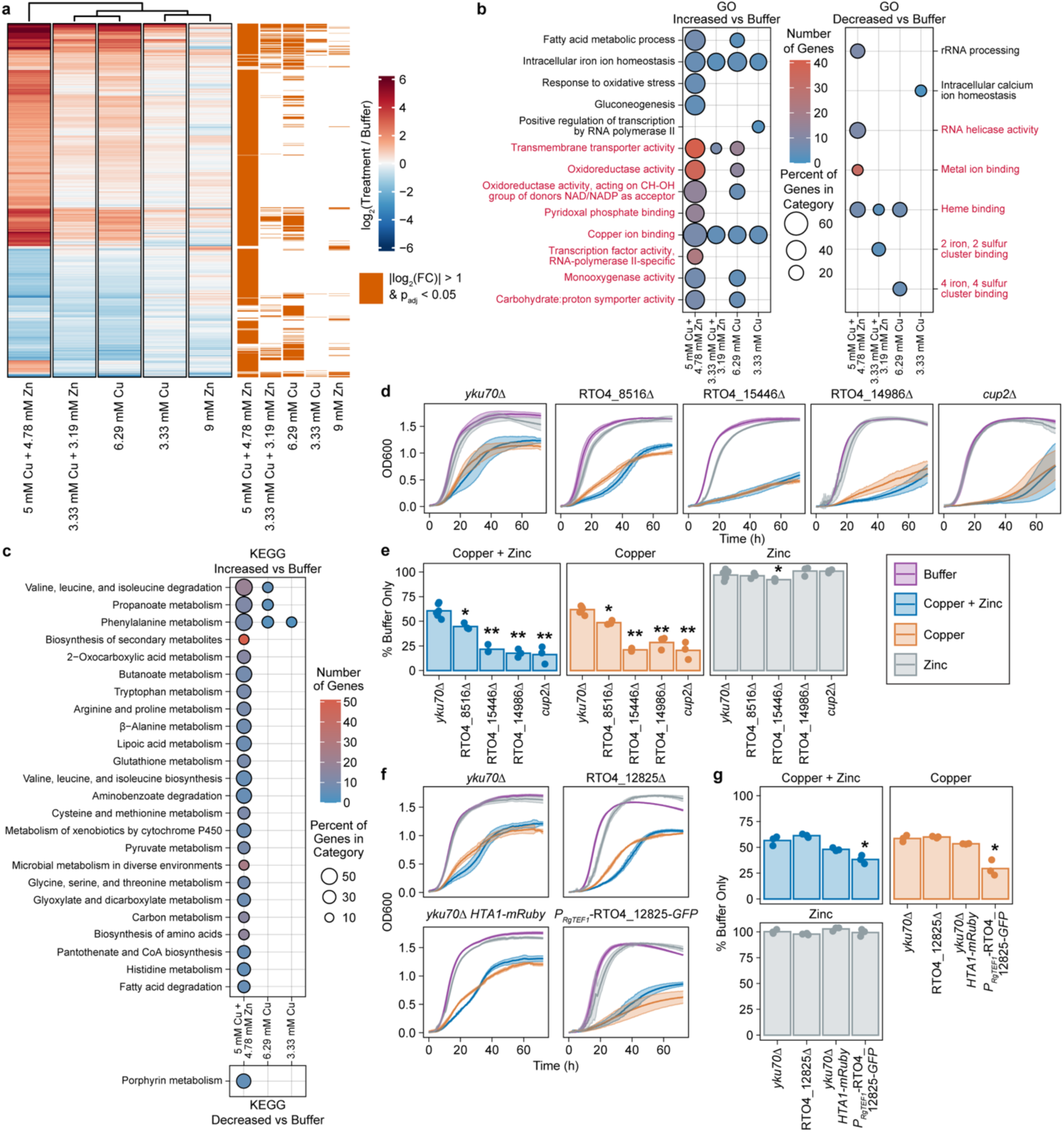
*R. toruloides* downregulates rRNA synthesis machinery and upregulates production of redox protective metabolites in response to copper and zinc. **A)** Heatmap of the log_2_(fold change) in gene expression during exposure to the indicated concentrations of copper sulfate and/or zinc sulfate relative to exposure to ammonium-phosphate buffer. Only genes at least two-fold differentially expressed in at least one condition are displayed. Orange bars to the right of the heatmap indicate genes at least two-fold differentially expressed in the indicated condition relative to buffer with p_adj_ < 0.05. RNAseq was performed with three biological replicates. **B-C)** (**B**) GO^34,35^ and (**C**) KEGG^36–38^ functional category enrichment analyses of genes at least two-fold upregulated ((**B**) left/(**C**) top) or downregulated ((**B**) right/(**C**) bottom) in the indicated conditions relative to buffer. Displayed comparisons had at least one significantly enriched category (p_adj_ < 0.05). Dot color and size represent the number of differentially expressed genes and percent of differentially expressed genes in the category, respectively. Text of GO biological process and molecular function categories is black and red, respectively. **D and F)** Growth (OD600) of the indicated deletion and *P_RgTEF1_-*RTO4_12825-*GFP* mutants and their parental strains, *yku70*Δ and *yku70*Δ *HTA1-mRuby*, respectively, on media supplemented with ammonium-phosphate buffer or ammonium-phosphate buffer with 7.5 mM copper sulfate, 9.56 mM zinc sulfate, or 5 mM copper sulfate plus 4.78 mM zinc sulfate. Lines are averages and colored bands are standard deviations of at least three biological replicates. **E and G)** Area under the curve from (**D**) for (**E**) or (**F**) for (**G**) of the indicated strains grown in the indicated conditions normalized to the area under the curve of the same strain on media supplemented with ammonium-phosphate buffer. Bars are averages and dots are individual data points of at least three biological replicates. *p < 0.05 and **p < 10^-9^ as determined by (**E**) one-way ANOVA with a Dunnett’s post-hoc test compared to the *yku70*Δ parental strain and (**G**) two-sample *t*-tests of RTO4_12825Δ and *P_RgTEF1_-*RTO4_12825-*GFP* cells compared to their respective parental strain. Statistical tests in (**E**) were performed using the growth of all strains assessed in Fig S9.

Functional enrichment analysis of genes increased in expression relative to buffer suggested that during exposure to copper and copper plus zinc, *R. toruloides* increased catabolism of fatty acids and branched-chain amino acids as anaplerotic substrates, presumably to enable increased production of the protective molecules cysteine and glutathione (Fig 4B-4C and S8). In 5 mM copper plus 4.78 mM zinc, most genes encoding mitochondrial fatty acid catabolism enzymes were upregulated, including 2.4, 3.9, and 8-fold increases in the genes encoding enzymes necessary for the rate limiting acylation and translocation steps via carnitine palmitoyltransferase I (RTO4_14245), carnitine:acylcarnitine translocase (RTO4_12518), and carnitine palmitoyltransferase II (RTO4_13580), respectively (Fig S8). Similarly, expression of genes predicted to encode branched-chain amino acid catabolic enzymes was increased, including branched-chain amino acid aminotransferases (RTO4_14014, RTO4_14610, and RTO4_14853) and four of five genes encoding the branched-chain α-keto acid dehydrogenase complex (RTO4_12566, RTO4_15436, RTO4_11188, and RTO4_11183) (Fig S8).

Fatty acid and branched-chain amino acid catabolism produce acetyl-CoA and propionyl-CoA, which can be processed into anaplerotic compounds via the glyoxylate and methylcitrate cycles, respectively. We observed a 77 and 16-fold upregulation of genes encoding the glyoxylate cycle enzymes isocitrate lyase (RTO4_14022) and malate synthase (RTO4_9457), respectively, and a 3.9 and 2.4-fold increase in expression of genes encoding the methylcitrate cycle enzymes 2-methylcitrate dehydratase (RTO4_12642) and methylisocitrate lyase (RTO4_14162) (Fig S8). Further, the anaplerotic signature gene pyruvate carboxylase (RTO4_16630) was upregulated 2.4-fold in 5 mM copper plus 4.78 mM zinc (Fig. S8).

Genes that may play a role in mitigating metal toxicity were upregulated on 5 mM copper plus 4.78 mM zinc, such as cysteine and glutathione synthesis genes. Synthesis of cysteine initiates from the tricarboxylic acid cycle with conversion of oxaloacetate to aspartate or transfer of a succinyl group from succinyl-CoA to serine. Key genes of both pathways were upregulated including a 29-fold increase of RTO4_9499, encoding cystathionine γ-lyase, and a 2.7-fold increase in RTO4_15248, encoding serine O-succinyltransferase. Expression of glutamate-cysteine ligase (RTO4_12007), the rate limiting enzyme in glutathione synthesis, and five glutathione-S-transferase encoding genes (RTO4_8431, RTO4_10132, RTO4_10933, RTO4_11814, and RTO4_16750) was also increased (Fig S8 and Datasets S3-S4). Additionally, several ferric reductases and multicopper oxidase genes were upregulated, which may decrease metal toxicity through alterations of metal oxidation state, as Cu^+^ is considered more toxic than Cu^2+^ (Fig 4B and Datasets S3-S4)^8^. Unlike *N. crassa*, the functions of genes downregulated in 5 mM copper plus 4.78 mM zinc were enriched for rRNA processing and RNA helicase activity (Fig 4B and Dataset S3-S4). Collectively, these data suggest *R. toruloides* increases anaplerosis and cysteine/glutathione biosynthesis while decreasing rRNA processing, and thereby likely ribosome biogenesis and/or protein synthesis, in response to copper and zinc.

To gain further insights into the genetic mechanisms underlying copper and zinc resistance we constructed deletion mutants of seven genes that were upregulated in multiple treatment conditions in *R. toruloides* (Fig 4D-4E, S9A-S9B, and Dataset S4). All deletion mutants grew normally with buffer or zinc treatment. However, deletion of RTO4_14231, encoding a copper-fist transcription factor homologous to *Saccharomyces cerevisiae* Cup2; the putative siderophore transporter RTO4_8516; the predicted heat shock transcription factor RTO4_14986; and the hypothetical protein RTO4_15446 increased sensitivity to both copper and copper plus zinc treatments relative to wild type cells (Fig 4D). Although RTO4_11264, RTO4_11240, and RTO4_16245 were over 40-fold upregulated on 5 mM copper plus 4.78 mM zinc, we did not see altered resistance to copper, zinc, or copper plus zinc treatment when these genes were deleted (Fig S9A-S9B and Dataset S4).

The most downregulated gene across both copper plus zinc and zinc only treatments was RTO4_12825, encoding a putative iron/zinc transporter homologous to Zrt1 in *S. cerevisiae* (Dataset S4). We hypothesized overexpression of RTO4_12825 would make *R. toruloides* sensitive to zinc. To test this, we generated an RTO4_12825Δ strain and an RTO4_12825 overexpression strain by addition of a second copy of the RTO4_12825 open reading frame with a C-terminal GFP-tag under control of the *Rhodotorula glutinis TEF1* promoter (*P_RgTEF1_-*RTO4_12825-*GFP*) and assessed growth (Fig S9C). Surprisingly, the *P_RgTEF1_-*RTO4_12825-*GFP* strain displayed increased sensitivity only to the copper plus zinc and copper only conditions (Fig 4F-4G). The metal specificity of RTO4_12825 cannot be explained due to differences in localization, since RTO4_12825-GFP appeared constitutively localized to the endoplasmic reticulum, similar to *S. cerevisiae* Zrt1^41–44^ (Fig S9D).

### High-throughput screens identify roles for genes involved in vacuolar trafficking and metal chelation in metal toxicity resistance

Given the physiological synergy exhibited in the *R. toruloides* copper and zinc response, we hypothesized *R. toruloides* would also exhibit synergy in the copper and zinc response at the genetic level. To test this, we utilized a barcoded insertional mutant library^26^. We grew the library in minimal media or minimal media supplemented with either buffer, 3.33 mM copper plus 3.18 mM zinc, 6.29 mM copper, or 9 mM zinc and calculated gene fitness by measuring the log_2_(fold change) in the relative abundance of all barcodes associated with insertions in a particular gene in the final population compared to the starting population population^26^. To identify genes with metal-specific phenotypes as opposed to those with pleiotropic phenotypes, we also computed fitness in the final population exposed to copper, zinc, or copper plus zinc relative to the buffer only control condition. We focused on genes that had a fitness greater than 2 or less than -2 and p_adj_ < 0.05 in the condition of interest relative to both the starting population and the buffer condition (Fig 5A-5C and S10 and Dataset S5). We identified only one gene (RTO4_12312) with a zinc fitness phenotype (Fig 5C). In contrast, 32 and 39 genes had copper plus zinc or copper fitness phenotypes, respectively (Fig 5A-5B and Dataset S5). If copper and zinc genetically synergized, we would expect a unique repertoire of genes with copper plus zinc-specific fitness phenotypes, and/or potentiation in the magnitude of gene fitness changes in the dual-treated compared to the individual treatments. However, fitness on copper alone and on copper plus zinc was highly correlated – 25 genes had fitness phenotypes on both copper and copper plus zinc and most gene fitness was of similar magnitude in the two conditions – providing no clear evidence of genetic synergy (Fig S10B and Dataset S5).

**Fig 5.**
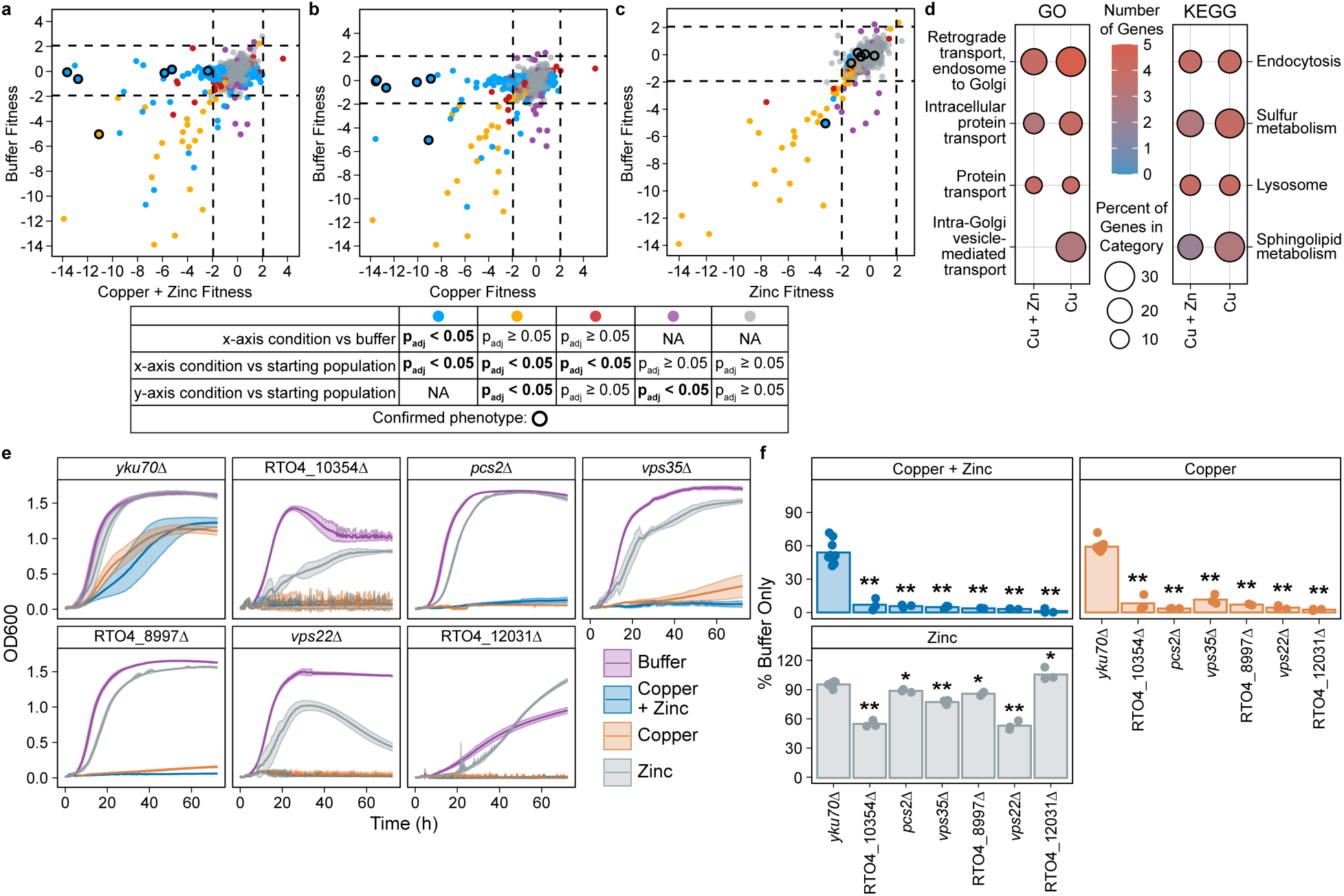
High-throughput screens identify roles for vacuolar trafficking and metal chelation in metal resistance in *R. toruloides*. **A-C)** Gene fitness scores^26^ relative to the starting population on media supplemented with ammonium-phosphate buffer or ammonium-phosphate buffer and **(A)** 3.33 mM copper plus 3.19 mM zinc, **(B)** 6.29 mM copper, or **(C)** 9 mM zinc. Genes with significant fitness scores (p_adj_ < 0.05) in the x-axis condition relative to the buffer condition and starting population are indicated in blue; in the x-axis and y-axis conditions relative to the starting population are indicated in yellow; in the x-axis condition relative to only the starting population are indicated in red; in the y-axis condition relative to the starting population are indicated in purple. Genes without a significant fitness score in the x-axis or y-axis condition relative to the starting population (p_adj_ ≥ 0.05) are indicated in grey. Genes with phenotypes confirmed by a clean deletion strain are circled in black. Fitness assays were performed with three biological replicates. **D)** GO^34,35^ (left) and KEGG^36–38^ (right) functional category enrichment analysis for genes with a significant (p_adj_ < 0.05) fitness less than -2 on 3.33 mM copper plus 3.19 mM zinc or 6.29 mM copper relative to the starting population and the ammonium-phosphate buffer. Dot color indicates the number of genes in the category and dot size indicates the percent of genes in the category that meet these fitness criteria. **E)** Growth (OD600) of the *yku70*Δ parental strain and indicated deletion mutant strains on media supplemented with ammonium-phosphate buffer or ammonium-phosphate buffer with 7.5 mM copper, 9.56 mM zinc, or 5 mM copper plus 4.78 mM zinc. **F)** Area under the curve from (**E**) normalized to the area under the curve of the same strain grown on media supplemented with ammonium-phosphate buffer. (**E**) Lines or (**F**) bars are averages and (**E**) colored bands are the standard deviations or (**F**) dots are individual data points of at least three biological replicates. *p < 0.05 and **p < 10^-9^ by one-way ANOVA with a Dunnett’s post-hoc test compared to the *yku70*Δ parental strain.

To examine the function of genes necessary for copper plus zinc and copper only resistance, we performed GO and KEGG enrichment analyses. Genes with negative copper or copper plus zinc fitness phenotypes were enriched for protein transport, endocytosis, sulfur metabolism, and sphingolipid metabolism (Fig 5D). Fungi combat metal toxicity by sequestering metals in the vacuole^45–48^. Most genes with decreased fitness in copper and copper plus zinc had roles in vacuole formation and trafficking, particularly via the ESCRT and AP-3 pathways (Dataset S5). These genes included two of the three genes encoding ESCRT-II complex proteins, (*VPS22/SNF8* [RTO4_9066] and *VPS36* [RTO4_10649]), *VTA1* (RTO4_14056) of the Vps4-Vta1 complex, the ARF-binding protein encoding gene *GGA1* (RTO4_9655), *VPS11* (RTO4_13845) of the CORVET complex, and the AP-3 complex delta subunit *APL5* (RTO4_15799). Genes encoding members of the Golgi-associated retrograde protein (GARP) (*VPS54* [RTO4_15239] and *VPS52* [RTO4_12310]) and retromer (*VPS35* [RTO4_10531]) complexes, critical for retrograde transport, were required for resistance to copper and copper plus zinc. Several genes encoding SNARE proteins were decreased in fitness in copper and copper plus zinc: *VTI1* (RTO4_9111), a Vti1-like protein RTO4_15699, a Vam7-like protein RTO4_8997, and a Bet1-like protein RTO4_9409 (Dataset 5).

We generated deletion mutants of six genes with decreased fitness scores in our high-throughput screens. These genes included: three genes with predicted roles in protein trafficking and vacuole formation (*VPS22/SNF8*, *VPS35*, and the Vam7-like SNARE RTO4_8997); RTO4_10354, encoding a glycosyltransferase; *PCS2* (RTO4_10995), encoding a phytochelatin synthase; and *MCY1* (RTO4_12031), encoding a cysteine synthase. Although *MCY1* had reduced fitness in buffer, we chose to delete it to determine the impact of blocking cysteine synthesis via serine while leaving cysteine synthesis via aspartate intact.

Three of the clean deletion mutants had growth defects on buffer. Cells lacking *MCY1* grew substantially worse than *yku70*Δ cells, while RTO4_10354Δ, and *vps22Δ* cells grew at a similar rate as *yku70*Δ cells but reached a lower final OD600 (Fig 5E). We exposed cells to 7.5 mM copper or 5 mM copper plus 4.78 mM zinc, which reduced growth of *yku70*Δ by 50% (Fig 5F). Corroborating our high-throughput screen, all deletion mutants had over 88% growth inhibition on 5 mM copper plus 4.78 mM zinc and 7.5 mM copper (Fig 5E-5F). Despite lacking a phenotype on zinc in our high-throughput screen, RTO4_10354Δ, *vps35*Δ, and *vps22*Δ cells also exhibited sensitivity to 9.56 mM zinc (Fig 5E-5F). These data suggest the glycosyltransferase RTO4_10354 plays a role in responding to metal toxicity and confirms roles for metal chelation and vacuolar function in metal toxicity resistance.

### RNAseq and high-throughput screens identify different genetic responses to copper and zinc

Transcriptional profiling and fitness screens of *R. toruloides* both identified genes necessary for resistance to copper and zinc toxicity (Fig 4 and 5). Although previous comparisons of gene expression with gene fitness have had mixed results^49–54^, we hypothesized transcriptional and genetic responses to copper and zinc toxicity would be partially correlated due to our identification of genes that are both transcriptionally regulated and required for resistance to copper and zinc (Fig 4). However, we did not observe any signs of correlation between fitness and expression changes, indicating the RNAseq and functional genomics datasets identified almost entirely distinct sets of metal toxicity response genes (Fig 6A-6C).

**Fig 6.**
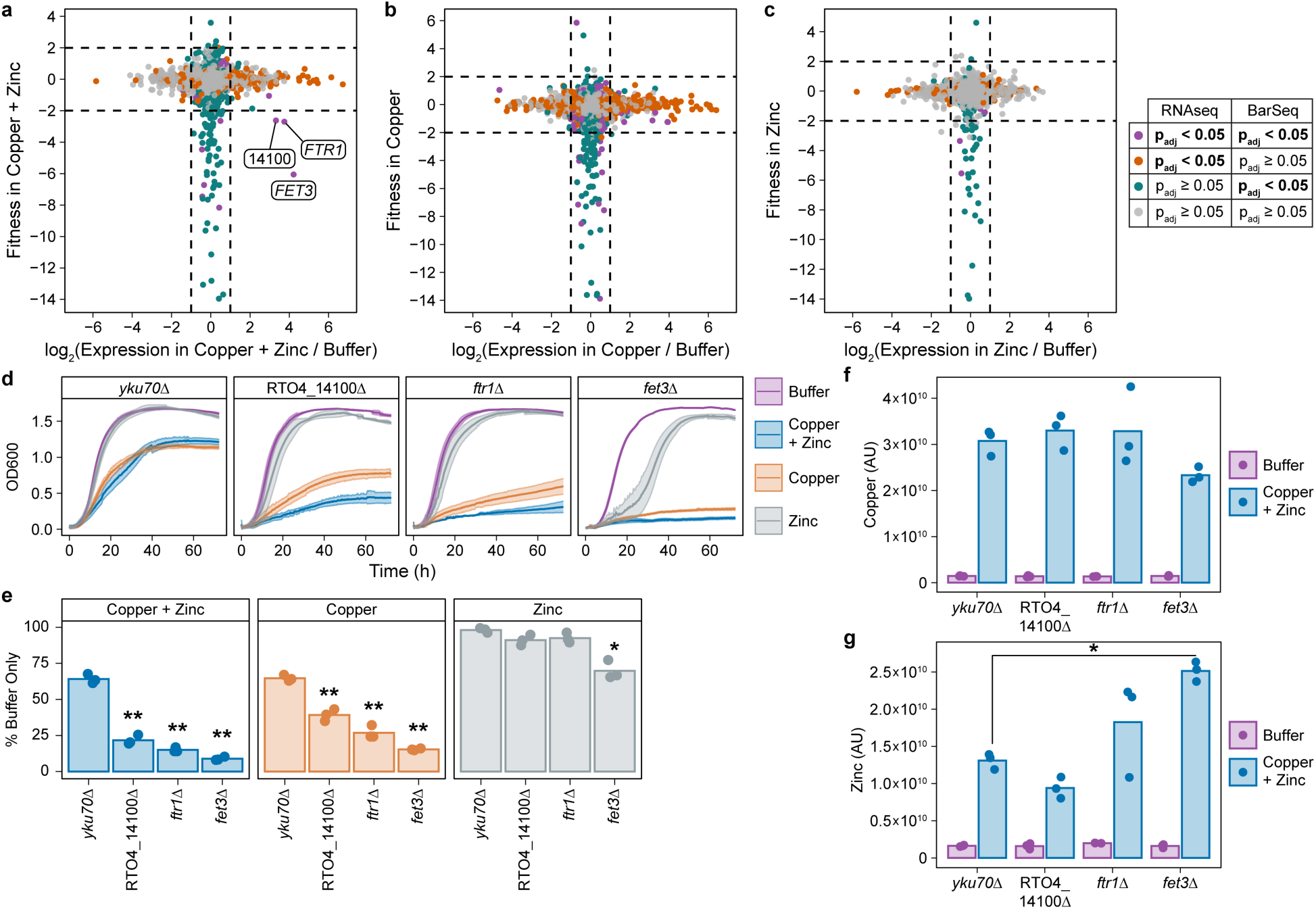
Fitness assays and transcriptomics identify different sets of genes involved in resistance to copper and zinc. **A-C)** Scatterplots of the log_2_(fold change) in *R. toruloides* gene expression during exposure to the indicated conditions relative to ammonium-phosphate buffer from RNAseq compared to gene fitness relative to the starting population from fitness assays (BarSeq) for (**A**) 3.33 mM copper sulfate plus 3.18 mM zinc sulfate, (**B**) 6.3 mM copper sulfate, and (**C**) 9 mM zinc sulfate. Purple indicates genes with p_adj_ < 0.05 in both the RNAseq and BarSeq analyses, orange indicates p_adj_ < 0.05 in the RNAseq and p_adj_ ≥ 0.05 in the BarSeq analyses, green indicates p_adj_ ≥ 0.05 in the RNAseq and p_adj_ < 0.05 in the BarSeq analyses, and grey indicates p_adj_ ≥ 0.05 in both the RNAseq and BarSeq analyses. Dotted black lines indicate a fitness score greater than 2 or less than -2 and differential expression of at least 2-fold. Genes significant in both the RNAseq (at least 2-fold differentially expressed) and BarSeq (fitness of greater than 2 or less than -2) analyses are labeled with their gene IDs. **D)** Growth (OD600) of the *yku70*Δ parental strain and indicated deletion mutants on media supplemented with ammonium-phosphate buffer or ammonium-phosphate buffer with 7.5 mM copper sulfate, 9.56 mM zinc sulfate, or 5 mM copper sulfate plus 4.78 mM zinc sulfate. **E)** Area under the curve from (**D**) of the indicated cells grown in the indicated conditions normalized to the area under the curve of the same strain grown on media supplemented with ammonium-phosphate buffer. **F-G)** Relative cellular concentrations of (**F**) copper and (**G**) zinc quantified using x-ray fluorescence imaging on an equal amount of the indicated *R. toruloides* cells grown in media supplemented with ammonium-phosphate buffer or ammonium-phosphate buffer with 5 mM copper and 4.78 mM zinc. (**D**) Lines or (**E-G**) bars are averages and (**D**) colored bands are the standard deviations or (**E-G**) dots are individual data points of at least three biological replicates. *p < 0.01 and **p < 10^-5^ by (**E**) one-way ANOVA with a Dunnett’s post-hoc test relative to the *yku70*Δ parental strain or (**F-G**) two-way ANOVA with a Tukey’s post-hoc test of the indicated mutants relative to *yku70*Δ cells during exposure to copper plus zinc.

Across all comparisons, only three genes were both differentially expressed by at least two-fold and had a fitness less than -2, and only in the copper plus zinc treated condition: RTO4_14100, RTO4_15844, and RTO4_9364 (Fig 6A). RTO4_14100 encodes a bZIP transcription factor with some homology to the *Cryptococcus neoformans* iron regulator HapX (24.9% identity and 35.2% similarity by EMBOSS Needle alignment^55^) but little to no homology to characterized Ascomycete HapX proteins. RTO4_15844 and RTO4_9364 are homologous to the *S. cerevisiae* high-affinity iron transporter Ftr1 and multicopper oxidase Fet3, respectively. In *S. cerevisiae*, Fet3 complexes with Ftr1 and oxidizes Fe^2+^ to Fe^3+^ whereupon Ftr1 transports Fe^3+^ into the cell^56–58^. Deletion of RTO4_14100, *FTR1* (RTO4_15844), or *FET3* (RTO4_9364) resulted in increased sensitivity to copper plus zinc and copper only treatments relative to *yku70*Δ cells (Fig 6D-6E). Additionally, *fet3*Δ cells were more sensitive to zinc treatment than *yku70*Δ cells, suggesting a conserved role for Fet3 in metal resistance in *R. toruloides* (Fig 6D-6E)^59–61^.

Because copper and zinc in combination changed the cellular quantity of each metal compared to treatment with each metal individually (Fig 2), we hypothesized the increased sensitivity of RTO4_14100Δ, *ftr1*Δ, and *fet3*Δ cells could be explained by alterations in cellular copper and/or zinc concentrations. To test this hypothesis, we performed x-ray fluorescence imaging on cells incubated in media supplemented with buffer alone or 5 mM copper plus 4.78 mM zinc. Copper and zinc concentrations in RTO4_14100Δ cells and copper concentrations in *ftr1*Δ and *fet3*Δ cells were indistinguishable from those in *yku70*Δ cells (Fig 6F and S2). However, zinc concentrations in *fet3*Δ cells were 92% higher than in *yku70*Δ cells, while zinc concentrations in *ftr1*Δ cells were highly variable (Fig 6G and S2). Thus, the increased sensitivity of *fet3*Δ cells to copper plus zinc may be due to increased cellular zinc concentrations.

The identification of a transcription factor, RTO4_14100, by both transcriptomics and fitness assays led us to hypothesize that RTO4_14100 may regulate transcription of metal-responsive genes. To test this hypothesis, we incubated *yku70*Δ and RTO4_14100Δ cells in media with buffer or copper plus zinc and performed reverse transcription quantitative PCR on five genes with increased expression on copper plus zinc (RTO4_8516, RTO4_11264, RTO4_12325, *FTR1*, and *FET3*) and one gene with decreased expression on copper plus zinc (RTO4_16473) (Datasets S3-S4). The predicted siderophore transporter RTO4_8516 and the hypothetical protein RTO4_16473 were both at least six-fold downregulated and the predicted dienelactone hydrolase RTO4_11264 was six-fold upregulated in copper plus zinc in RTO4_14100Δ cells relative to *yku70*Δ cells, demonstrating that RTO4_14100 directly or indirectly regulates genes in response to metal exposure (Fig S11).

## DISCUSSION

The beneficial impact of copper and zinc in combination was first noted in 1942 as an observation that Florida tomato crops performed better when zinc sulfate was added to a copper fungicide^62^. However, the reason behind this observation was never determined. More recent studies show potential synergy of copper and zinc toxicity in the bacteria *Vibrio fischeri*^63^ and *Escherichia coli*^18^ and the nematode *Caenorhabditis elegans*^64^. We demonstrate copper and zinc interact synergistically to inhibit both Ascomycete and Basidiomycete fungal growth (Fig 1 and S1), likely due, at least partially, to increased cellular zinc when copper is present (Fig 2B). Zinc stress inhibits transport of other ions^17,18^. Correspondingly, cellular copper was decreased when cells were exposed to copper plus zinc (Fig 2A). A small number of genes exhibited transcriptional synergy in the *N. crassa* response to copper plus zinc, potentially due to increased cellular zinc concentrations, the application of two metals with similar modes of action^10,13,14,16,19–21^, or both. Long-term copper fungicide use leads to copper accumulation in soils^65^. Our data suggest copper and zinc combination fungicides, like Curezin^®^, require lower copper application levels for effective fungal inhibition, increasing pest management sustainability, particularly in organic agriculture where copper is one of the few fungicides approved for use^66^.

Although both *N. crassa* and *R. toruloides* upregulated expression of cysteine and glutathione biosynthesis in response to copper and zinc (Fig 3 and 4), likely to protect against oxidative stress and inhibit Fenton-like reactions^8,12^, the *N. crassa and R. toruloides* transcriptional responses to copper and zinc otherwise differed substantially. *N. crassa* broadly decreased its metabolism in response to copper and zinc. In contrast, *R. toruloides* shifted its metabolism towards anaplerosis via catabolism of fatty acids and branched chain amino acids, corroborating previous transcriptomic and proteomic studies of the *Rhodotorula*/*Rhodosporidium* response to metal toxicity^45,67–69^. *N. crassa* increased expression of protein stabilizing machinery, such as heat shock proteins, and genes associated with rRNA and ribosome biogenesis in response to copper and copper plus zinc stress, likely to protect against proteotoxic stress from protein mismetallation, denaturation, and aggregation^8,9,16,18^ (Fig 3A-3C). Surprisingly, we observed no such increase in heat shock protein expression and a downregulation of genes associated with rRNA synthesis and mitochondrial ribosomal proteins in *R. toruloides* (Fig 4A-4C). Differences in roles for rRNA and ribosomal genes in metal toxicity responses between *N. crassa* and *R. toruloides* may be indicative of differences present across Ascomycete and Basidiomycete fungi, as ribosome biogenesis genes were upregulated in response to copper in the ascomycete *Candida albicans* and ribosome biogenesis proteins were downregulated in response to copper in the basidiomycete *C. neoformans*^70,71^.

Genetic characterization of genes involved in metal responses can help to predict future mechanisms of fungicide resistance. Loss-of-function mutations causing increased metal resistance are particularly concerning, given the multitude of mechanisms through which loss-of-function mutations can evolve. Deletion of three genes increased resistance to zinc in *N. crassa*: the pyruvate decarboxylase *cfp*; the transcription factor *znf-27*; and the heat shock factor protein *hsp30*, which was particularly surprising given heat shock factors’ roles in protein stabilization and stress resistance (Fig 3D-3F). Critically, these loss-of-function mutations did not increase resistance to copper and zinc in combination, providing an additional advantage to utilizing this metal combination to treat fungal disease.

We identified genes necessary for resistance to copper and zinc toxicity using both transcriptional profiling and high-throughput fitness assays in *R. toruloides*. Despite this, there was essentially no overlap between differentially expressed genes and genes with fitness phenotypes (Fig 6). This lack of correlation has been observed in several bacteria and *S. cerevisiae*^49,51,53,72,73^ and is likely due, at least in part, to genes that play a role but are not transcriptionally regulated in the tested condition, or genes that are transcriptionally regulated but have a non-critical or redundant role in the tested condition. Perhaps most surprisingly, we identified two genes using transcriptomics that were required to survive metal toxicity but did not have a phenotype in our pooled fitness assays: RTO4_8516 and RTO4_14986. One possible mechanism for this phenomenon is that mutations in genes associated with secreted proteins/molecules are complemented in pooled assays due to the availability of secreted public goods. This mechanism could explain why the predicted siderophore transporter RTO4_8516 loss-of-function mutants did not exhibit a phenotype in pooled fitness assays and may give hints into the molecular mechanism of RTO4_14986 in the metal response.

Prior use of high-throughput screens with transcriptomics in *R. toruloides* demonstrated genes with both reduced fitness and increased expression in a condition likely play a role in responding to that condition^50^. We identified three genes with fitness defects and increased expression on copper plus zinc and demonstrated clear roles for these genes in copper and copper plus zinc resistance. The transcription factor RTO4_14100 directly or indirectly regulates expression of several metal-responsive genes, including the siderophore transporter gene RTO4_8516, which plays a role in the copper response (Fig 4D-4E and Fig S11). *FET3* and *FTR1* are well known mitigators of copper toxicity in *S. cerevisiae*^56–59,61^, where Fet3 oxidizes Cu^+^ to the less toxic Cu^2+^ form and Ftr1 is necessary for proper localization and protein levels of Fet3^56,58,59^. *S. cerevisiae* cells lacking *FET3* have increased cellular copper^61^. In contrast, *R. toruloides* cells lacking *FET3* have increased zinc, but not copper, following exposure to copper plus zinc (Fig 6F-6G). Taken together, our study demonstrates the critical complementarity of transcriptomics with high-throughput screens in identifying genetic mechanisms of stress or antimicrobial resistance. We expect our findings to inform future investigations of genetic mechanisms of fungal responses to fungicides and antifungal drugs.

## MATERIALS and METHODS

### Strains and culture conditions

*N. crassa* strains used in this study are listed in Table S1. Strains were derived from the wild type strain, FGSC 2489, and obtained from the *N. crassa* deletion collection^74,75^. Both the wild type *mat A* strain (FGSC 2489) and the wild type *mat a* strain (FGSC 4200) were used as wild type controls for initial mutant screening (Fig S5). FGSC 2489, the wild type *mat A* strain, was used as the wild type strain for all other experiments. *N. crassa* cells were grown from 25% glycerol freezer stocks on Vogel’s minimal medium^27^ + 2% sucrose + 1.5% agar (BD Difco^TM^ Bacto^TM^) slants for 2 d at 28°C in the dark and 4 d at 28°C in constant light. *N. crassa* conidia were harvested from slants with sterile double distilled H_2_O for inoculation. All *N. crassa* experiments were performed using filter-sterilized Vogel’s minimal medium^27^ with 50 mM NH_4_Cl replacing NH_4_NO_3_ as the nitrogen source (VMM NH_4_Cl) and 2% w/v glucose as the carbon source.

*R. toruloides* strains used in this study are listed in Table S2. Strains were derived from the wild type reference strain IFO 0880 (also called NBRC 0880, obtained from the Biological Resource Center, NITE (NBRC), Japan). Wildtype IFO 0880 is the background strain for the previously constructed barcoded insertional mutagenesis library^26^ used in this study (Fig 5A-C and S10) and was used for the experiments characterizing copper and zinc interactions (Fig 1C) and RNAseq (Fig 4A-C). The background strain for all genetic manipulations was the non-homologous end-joining deficient *yku70Δ* (RTO4_11920Δ) strain^76^. *R. toruloides* strains were struck for singles on yeast peptone dextrose (YPD) (VWR) + 2% agar (BD Difco^TM^ Bacto^TM^) from 25% glycerol freezer stocks and grown at 28°C for 2 d to 3 d. Single colonies were inoculated into liquid YPD and grown at 28°C shaking at 250 rpm to exponential phase prior to starting experiments. All *R. toruloides* experiments were performed using filter-sterilized VMM NH_4_Cl with 2% w/v glucose as the carbon source.

### Copper sulfate and zinc sulfate stocks and media preparation

The ammonium-phosphate buffer was generated by dissolving 62.5 g/L phosphoric acid in water and bringing the solution to pH 2 using ammonium hydroxide. Copper sulfate pentahydrate (Fisher) and zinc sulfate heptahydrate (J. T. Baker) were dissolved in the ammonium-phosphate buffer at concentrations of 1.2 M and 1.53 M, respectively. Fresh copper sulfate stocks were made every one to two weeks to avoid formation of precipitates. Copper sulfate and zinc sulfate stocks were added to filter-sterilized VMM NH_4_Cl + 2% glucose at the indicated concentrations for each experiment. To control for the low pH of the buffer, addition of phosphate and ammonium as potential nutrients, and minor alteration of the concentration of VMM NH_4_Cl + 2% glucose, ammonium-phosphate buffer was added to each condition, unless specified, such that the total volume of all media supplementations were equal across all conditions within an experiment.

### R. toruloides and N. crassa growth experiments

To measure *N. crassa* growth, *N. crassa* conidia were inoculated at 10^6^ conidia/mL into either 100 mL of liquid media in 250 mL flasks or 3 mL of liquid media in deep-well 24-well plates (initial mutant screen in Fig S5 only). Cultures were grown for 48 h in constant light at 28°C with constant shaking at 200 rpm. Biomass was harvested via vacuum filtration, washed with double distilled H_2_O, dried for 24 h in a 60°C drying oven, and weighed to determine biomass accumulation. Assessment of *N. crassa* mutant growth was performed in VMM NH_4_Cl + 2% glucose supplemented with ammonium-phosphate buffer or ammonium-phosphate buffer with 3.33 mM copper sulfate, 3.19 mM zinc sulfate, or 2 mM copper sulfate plus 1.91 mM zinc sulfate.

To measure *R. toruloides* growth, colonies of *R. toruloides* were grown in liquid YPD at 28°C with constant shaking at 250 rpm until exponential phase. Cells were pelleted at 4000 g and washed three times with VMM NH_4_Cl + 2% glucose. Washed cells were used to inoculate 750 µL of media in 48-well plates at a starting OD600 of 0.1, as measured by an Eppendorf BioPhotometer with a 10 mm path length. Growth assays were conducted using an Agilent BioTek Epoch2 plate reader. Cultures were incubated at 28°C with constant shaking (3 mm orbit, 425 rpm). OD600 measurements were taken every 20 minutes for 48 h (characterization of copper and zinc interactions in wild type [Fig 1 and S1]) or 72 h (mutant characterization [Fig 4, 5, 6, and S9). Assessment of *R. toruloides* mutants was done in VMM NH_4_Cl + 2% glucose supplemented with ammonium-phosphate buffer or ammonium-phosphate buffer with 7.5 mM copper sulfate, 9.56 mM zinc sulfate, or 5 mM copper sulfate plus 4.78 mM zinc sulfate.

### Synergy calculation

Synergy was calculated using the R package SynergyFinder Plus v. 3.18.0^28^. Percent inhibition relative to an untreated control was calculated using biomass for *N. crassa* and area under the curve for *R. toruloides* and used as input for the CalculateSynergy function. Contour plots were generated using the Plot2DrugContour function.

### R. toruloides strain construction

Transformation of *R. toruloides* was carried out using *Agrobacterium tumefaciens* mediated transformation as previously described^26^ using *A. tumefaciens* EHA 105 and plasmids derived from pGI2^77^. Gene deletions were made in the *yku70*Δ background using homologous recombination of a nourseothricin resistance cassette with ∼1 kb homology arms up and downstream of the gene’s open reading frame. To generate the *HTA1-mRuby* strain, we fluorescently tagged the histone H2A subunit gene *HTA1* (RTO4_12090) with the mRuby2 fluorescent protein at the native locus. The *HTA1-mRuby* construct was integrated using a combination of G418 resistance and a 5-fluorodeoxyuridine (FUDR) sensitive thymidine kinase counterselection marker flanked by 250 bp terminal repeat sequences. The construct was transformed into the *yku70Δ* background using G418 to select for transformants. Transformed colonies were passages one time on liquid YPD and plated onto yeast nitrogen base with (NH_4_)SO_4_ without amino acids (MP Biomedicals) + 1% glucose + 50 µg/mL FUDR (VWR) to select for strains that looped out the G418 resistance and thymidine kinase markers. Colonies were screened by microscopy, PCR, and locus sequencing for correct integration of the mRuby2 tag and successful removal of the G418 and thymidine kinase markers. The RTO4_12825-*GFP* overexpression strain was generated in the *HTA1-mRuby* background. The open reading frame of RTO4_12825 was C-terminally tagged with meGFP and placed under control of the *TEF1* promoter from *Rhodotorula graminis* strain WP-1^79^ and the *T_nos_* terminator^80^. A construct with a G418 resistance cassette and *P_RgTEF1_-*RTO4_12825-*GFP*-*T_nos_* was targeted to the *YKU70* locus with ∼1 kb arms of homology up and downstream of the hygromycin resistance cassette used to initially generate the deletion of *YKU70*^76^. Nourseothricin (GoldBio) or G418 (VWR) were added to media at 100 µg/mL to select transformants. All gene deletion strains were confirmed by PCR for absence of the deleted gene and correct integration of the resistance cassette. The *P_RgTEF1_*-RTO4_12825-*GFP* strain was confirmed by PCR and DNA sequencing for correct locus integration and absence of mutations, and reverse-transcription quantitative PCR for baseline expression of RTO4_12825 compared to the *yku70*Δ *HTA1-mRuby* parental strain (Fig S9C).

### Copper and zinc quantification using x-ray fluorescence imaging

Colonies of *R. toruloides yku70*Δ, RTO4_14100Δ, *ftr1*Δ, and *fet3*Δ strains were grown to exponential phase in YPD at 28°C with shaking at 250 rpm. Cultures were pelleted at 4000 g and washed three times with VMM NH_4_Cl + 2% glucose. Washed cells were used to inoculate 100 mL of VMM NH_4_Cl + 2% glucose supplemented with ammonium-phosphate buffer or ammonium-phosphate buffer with 5 mM copper sulfate plus 4.78 mM zinc sulfate for all cells and also ammonium-phosphate buffer with 5 mM copper sulfate or 4.78 mM zinc sulfate for *yku70*Δ cells in 250 mL flasks at a starting OD600 of 0.1. Cultures were incubated for 18 h at 28°C with shaking 250 rpm. Twenty OD600 units of cells, as measured by an Eppendorf BioPhotometer with a 10 mm path length, were pelleted at 4000 g and washed three times with VMM NH_4_Cl + 2% glucose. Washed cells were resuspended in 1 mL of VMM NH_4_Cl + 2% glucose and quantified. Ten OD600 units of cells were transferred to an Eppendorf tube, pelleted at 7500 g, and resuspended in 10 µL of VMM NH_4_Cl + 2% glucose. Cells were spotted onto 12.7 µm Kapton film (American Durafilm) and dried onto the Kapton film at 65°C. An equal volume of VMM NH_4_Cl + 2% glucose supplemented with ammonium-phosphate buffer without any cells was also spotted onto the Kapton and dried as a control.

The x-ray fluorescence imaging data were collected using a single-element silicon drift diode Vortex detector (Hitachi, Japan) paired with an Xspress 3 mini digital signal processor (Quantum Detectors, UK). Incident x-rays were generated with an XTG UltraBright Microfocus Source with a tungsten anode (Oxford, UK) operated at 50 kV and 1.6 mA (maximum power 80 W), paired with a custom polycapillary optic (XOS, USA) to focus the beam size to approximately 200 μm, matching the step size used for data collection. X-ray fluorescence data were acquired in a continuous flyscanning mode with a dwell time of 800 ms per 200 µm bin across the two-dimensional map. Praxes v. 0.7.1 (https://github.com/praxes/praxes), a Cornell High Energy Synchrotron Source (CHESS)-developed Python interface wrapping PyMca^81^ libraries, was used to fit the x-ray fluorescence spectra, producing elemental distribution maps. All fungal samples were prepared with a consistent optical density, and all samples were measured within a single scan, allowing for direct comparison of the relative concentrations of Cu and Zn between samples.

### RNA sequencing and transcript abundance

To perform RNA sequencing (RNAseq) on *N. crassa*, 10^8^ wild type *mat A* conidia were inoculated into 100 mL of VMM NH_4_Cl + 2% glucose in 250 mL flasks and incubated at 28°C with shaking at 200 rpm for 16 h. The mycelial mass was then washed once in media containing the respective treatment condition (VMM NH_4_Cl + 2% glucose, VMM NH_4_Cl + 2% glucose + ammonium-phosphate buffer, VMM NH_4_Cl + 2% glucose + ammonium-phosphate buffer + 3.33 mM copper sulfate, VMM NH_4_Cl + 2% glucose + ammonium-phosphate buffer + 5 mM copper sulfate, VMM NH_4_Cl + 2% glucose + ammonium-phosphate buffer + 3.19 mM zinc sulfate, VMM NH_4_Cl + 2% glucose + ammonium-phosphate buffer + 4.78 mM zinc sulfate, VMM NH_4_Cl + 2% glucose + ammonium-phosphate buffer + 2 mM copper sulfate + 1.91 mM zinc sulfate, or VMM NH_4_Cl + 2% glucose + ammonium-phosphate buffer + 3.33 mM copper sulfate + 3.19 mM zinc sulfate) and transferred to a fresh flask containing 100 mL of VMM NH_4_Cl + 2% glucose, VMM NH_4_Cl + 2% glucose + ammonium-phosphate buffer, VMM NH_4_Cl + 2% glucose + ammonium-phosphate buffer + 3.33 mM copper sulfate, VMM NH_4_Cl + 2% glucose + ammonium-phosphate buffer + 5 mM copper sulfate, VMM NH_4_Cl + 2% glucose + ammonium-phosphate buffer + 3.19 mM zinc sulfate, VMM NH_4_Cl + 2% glucose + ammonium-phosphate buffer + 4.78 mM zinc sulfate, VMM NH_4_Cl + 2% glucose + ammonium-phosphate buffer + 2 mM copper sulfate + 1.91 mM zinc sulfate, or VMM NH_4_Cl + 2% glucose + ammonium-phosphate buffer + 3.33 mM copper sulfate + 3.19 mM zinc sulfate. Flasks were incubated at 28°C with shaking at 200 rpm for eight hours. Twenty-five mL of biomass was vacuum filtered onto a 70 mm Whatman filter paper No. 1 (Cytiva) and flash frozen in liquid nitrogen for RNA extraction.

*N. crassa* RNA was extracted using a TRIzol^TM^ (Invitrogen) extraction similar to previously described^82^. Briefly, one mL of TRIzol^TM^ (Invitrogen) was added to a tube containing frozen biomass and 0.5 mm zirconia beads (BioSpec) on ice. Tubes were bead beaten for 45 s, rested on ice for one minute, then bead beaten for another 45 s. Samples were incubated at room temperature for five minutes followed by addition of 200 µL of chloroform and mixing by inversion. Phases were separated by centrifugation at 4°C and RNA in the aqueous phase was precipitated using isopropanol. The RNA pellet was washed with 75% ethanol, dried, and resuspended in nuclease-free water. RNA was treated with DNase I (NEB) for 10 min at 37°C followed by purification using the Monarch® Spin RNA Cleanup Kit (NEB).

To perform RNAseq on *R. toruloides*, *R. toruloides* IFO 0880 wild type cells were grown in YPD at 28°C with shaking (250 rpm) to exponential phase. Cells were pelleted at 4000 g and washed three times with VMM NH_4_Cl + 2% glucose. Washed cells were inoculated at an OD600 of 0.1, as measured by an Eppendorf BioPhotometer with a 10 mm path length, into 100 mL of VMM NH_4_Cl + 2% glucose, VMM NH_4_Cl + 2% glucose + ammonium-phosphate buffer, VMM NH_4_Cl + 2% glucose + ammonium-phosphate buffer + 3.33 mM copper sulfate, VMM NH_4_Cl + 2% glucose + ammonium-phosphate buffer + 5 mM copper sulfate, VMM NH_4_Cl + 2% glucose + ammonium-phosphate buffer + 9 mM zinc sulfate, VMM NH_4_Cl + 2% glucose + ammonium-phosphate buffer + 3.33 mM copper sulfate + 3.19 mM zinc sulfate, or VMM NH_4_Cl + 2% glucose + ammonium-phosphate buffer + 5 mM copper sulfate + 4.78 mM zinc sulfate in 250 mL flasks. Flasks were incubated for eight hours at 28°C with shaking at 200 rpm. Cultures were pelleted at 4000 g, the supernatant was removed, and cell pellets were flash frozen in liquid nitrogen. RNA was extracted using the Quick-RNA Fungal/Bacterial Miniprep Kit (Zymo) according to the manufacturer’s instructions. RNA was treated with DNase I (NEB) for 10 min at 37°C and purified using the Monarch® Spin RNA Cleanup Kit (NEB).

RNA integrity was assessed for all samples via agarose gel electrophoresis and for a randomly chosen subset of samples as a quality spot-check using an Agilent BioAnalyzer at the Biotechnology Resource Center at Cornell University. RNA was quantified using the Qubit RNA BR assay kit with a Qubit 2.0 (Invitrogen) and submitted to the Biotechnology Resource Center at Cornell University for library preparation and sequencing.

Libraries were prepared using the Lexogen QuantSeq 3’ mRNA-Seq Library Prep Kit FWD V2, and RNA was sequenced on an Illumina NextSeq 500 with 75 bp single-end reads at a depth of approximately 7.5 million reads per sample. Raw reads were trimmed using fastp v. 0.23.4 with the --trim_poly_x setting^83^. Reads from *N. crassa* samples were aligned to the *N. crassa* OR74A genome (v 12)^84^, and reads from *R. toruloides* samples were aligned to the *R. toruloides* IFO 0880 genome (v 4)^26^ using STAR v. 2.7.11b^85^ with --alignIntronMin set to 5. BAM files were indexed using SAMtools v. 1.20^86^. GNU parallel v. 20170522^87^ was used to run HTSeq v. 2.0.9^88^ in parallel for read counting with --nonunique set to random and -i set to Parent. For *R. toruloides* a gff3 file with improved 3’UTR annotation from Reyes-Chavez, Kerkaert, and Huberman 2026^78^ was used for running HTSeq.

Differential expression was determined using DESeq2 v. 1.50.1^89^. Counts per million (CPM) values were determined by dividing the counts for a given gene by the sum of all counts for the sample and multiplying by 10^6^. Genes were designated as significantly differentially expressed if they had an average CPM greater than 5 in at least one of the two compared conditions, p_adj_ < 0.05, and log_2_(fold change) greater than 2 or less than -2 for *N. crassa* or a log_2_(fold change) greater than 1 or less than -1 for *R. toruloides*. The ComplexHeatmap R package was used to visualize log_2_(fold change) relative to buffer (Fig 3A and 4A)^90^. Euclidean distance and hierarchical clustering were performed on log_2_(fold change relative to buffer) values for all genes significantly differentially expressed in at least one condition relative to buffer (Fig 3A and 4A). Principal component analysis was performed on log_2_ transformed normalized counts, generated using the normTransform function of DESeq2^89^, of the 2000 genes with the highest variance across all conditions for *N. crassa* and 1000 genes with the highest variance across all conditions for *R. toruloides*. RNAseq data were deposited in the Gene Expression Omnibus (GEO) at the National Center for Biotechnology Information (NCBI) and are accessible through GEO series accession numbers GSE348969 (*N. crassa*) and GSE348884 (*R. toruloides*).

### Functional enrichment analysis

Gene Ontology (GO)^34,35^ and Kyoto Encyclopedia of Genes and Genomes (KEGG)^36,38^ enrichment analyses were performed using the enricher function of the clusterProfiler R package (v. 4.18.1) with Benjamini-Hochberg multiple hypothesis testing corrections^91^. Significant enrichments were visualized using the clusterProfiler dotplot function. *R. toruloides* GO annotations were obtained from UniProt (Proteome ID: UP000239560)^92^ and KEGG annotations were obtained from Reyez-Chavez, Kerkaert, and Huberman (2026)^78^. KEGG annotations for *R. toruloides* were mapped to KEGG pathways using the clusterProfiler gson_KEGG_mapper function^91^ and pathways not relevant to fungi were manually removed (e.g. insect hormone biosynthesis, olfactory transduction, etc.). *N. crassa* GO annotations were obtained from FungiDB (release 68)^93,94^ and KEGG annotations were obtained from the KEGG database (organism code: ncr)^36,38^. Genes from the RNAseq experiments were included if they met the following differential expression relative to buffer thresholds for each organism: |log_2_(fold change)| > 2 & p_adj_ < 0.05 for *N. crassa* and |log_2_(fold change)| > 1 & p_adj_ < 0.05 for *R. toruloides* (Fig 3B-C and 4B-C). Genes from the fitness assays (BarSeq) experiments were included if they had fitness values less than -2 and p_adj_ < 0.05 in the comparisons of the condition of interest to both the starting population and buffer condition (Fig 5D).

### Reverse transcription quantitative PCR

To quantify RTO4_12825 expression in the *P_RgTEF1_-RTO4_12825-GFP* strain, the *yku70*Δ *HTA1-mRuby* parental strain and *P_RgTEF1_-RTO4_12825-GFP* cells were grown to exponential phase in liquid YPD. Cells were pelleted at 4000 g and washed three times with VMM NH_4_Cl + 2% glucose. Washed cells were inoculated into 5 mL of VMM NH_4_Cl + 2% glucose in 14 mL culture tubes at an OD600 of 0.1, as measured by an Eppendorf BioPhotometer with a 10 mm path length, and incubated at 28°C with shaking at 250 rpm. After eight hours cells were pelleted at 4000 g, supernatants were removed, and cell pellets were flash frozen in liquid nitrogen for RNA extraction.

To quantify expression of copper and zinc regulated genes in the RTO4_14100Δ strain, RTO4_14100Δ and *yku70*Δ cells were grown to exponential phase in liquid YPD. Cells were pelleted at 4000 g and washed three times with VMM NH_4_Cl + 2% glucose. Washed cells were inoculated into 100 mL of VMM NH_4_Cl + 2% glucose supplemented with either ammonium-phosphate buffer or ammonium-phosphate buffer with 3.33 mM copper sulfate plus 3.19 mM zinc sulfate in 100 mL flasks at an OD600 of 0.1, as measured by an Eppendorf BioPhotometer with a 10 mm path length. Flasks were incubated at 28°C with shaking at 200 rpm. After eight hours cells were pelleted at 4000 g, supernatants were removed, and cell pellets were flash frozen in liquid nitrogen for RNA extraction.

RNA was extracted using the Quick-RNA Fungal/Bacterial Miniprep Kit (Zymo) according to the manufacturer’s instructions. Purified RNA was treated with DNase I (NEB) for 10 min at 37°C and purified using the Monarch® Spin RNA Cleanup Kit (NEB). RNA quality was assessed via agarose gel electrophoresis and quantified via 260 nm absorbance using a Take3 microvolume plate (Agilent) on an Agilent BioTek Synergy HTX plate reader. Reverse transcription quantitative PCR (RT-qPCR) was performed using the Luna^®^ Universal One-Step RT-qPCR Kit (NEB) according to manufacturer’s instructions with 50 ng RNA as template. Data were collected on a CFX96^TM^ Real-Time System (Bio-Rad). Gene expression was normalized to the nuclear pore protein 84 homolog RTO4_9189. RT-qPCR primers are listed in Table S3.

### Microscopy

*P_RgTEF1_-RTO4_12825-GFP* cells were grown to exponential phase in liquid YPD. Cells were pelleted at 4000 g and washed three times with VMM NH_4_Cl + 2% glucose. Washed cells were inoculated at an OD600 of 0.1, as measured by an Eppendorf BioPhotometer with a 10 mm path length, into 750 µL of VMM NH_4_Cl + 2% glucose supplemented with ammonium-phosphate buffer or ammonium-phosphate buffer with 7.5 mM copper sulfate, 9.56 mM zinc sulfate, or 5 mM copper sulfate plus 4.78 mM zinc sulfate in a 48-well plate. Cells were incubated for eight hours at 28°C with shaking (3 mm orbit, 600 rpm) followed by mixing with 700 µL of cells with 77.8 µl 37% fresh molecular grade formaldehyde (Fisher) (to reach a final concentration of 3.7%). Cells were fixed at room temperature for 15 min followed by pelleting at 7500 g. The supernatant was removed and the cells were washed with phosphate-buffered saline (PBS) prior to resuspension in 100 µL of PBS + 25 µg/mL calcofluor white. After 5 min of incubation, cells were imaged on agarose coated slides using a Nikon Eclipse Ti2-E inverted microscope equipped with a prime BSI express CMOS camera using a CFI Plan Apochromat 100X oil objective.

### Fitness assays and analysis

Three aliquots of the *R. toruloides* random barcoded insertional mutagenesis library generated by Coradetti et. al. (2018)^26^ were each inoculated from freezer stocks into 100 mL of YPD in 250 mL flasks and incubated at 30°C with shaking at 250 rpm for 7.6 h (corresponding to ∼2.5-3 generations). Cells were pelleted at 4000 g, supernatants were removed, and cells were washed two times in VMM NH_4_Cl + 2% glucose. Washed cells were used to inoculate 100 mL VMM NH_4_Cl + 2% glucose, VMM NH_4_Cl + 2% glucose supplemented with ammonium-phosphate buffer, or VMM NH_4_Cl + 2% glucose supplemented with ammonium-phosphate buffer with 6.29 mM copper sulfate, 9 mM zinc sulfate, or 3.33 mM copper sulfate plus 3.19 mM zinc sulfate in 250 mL flasks at an OD600 of 0.1, as measured by an Eppendorf BioPhotometer with a 10 mm path length. The remainder of the washed cells were harvested by centrifuging at 4000 g and flash freezing the cell pellet in liquid nitrogen for quantification of relative barcode abundance in the starting (T0) populations. Cultures were incubated at 30°C with shaking at 250 rpm for approximately five to seven generations (i.e., ∼20 h). Cells were harvested by centrifuging at 4000 g and flash freezing the cell pellet in liquid nitrogen. Genomic DNA was extracted using the Quick-DNA Fungal/Bacterial Miniprep Kit (Zymo Research).

Barcodes were amplified using 200 ng of gDNA with Q5 high-fidelity polymerase (NEB). The PCR mix contained GC enhancer, Q5 reaction buffer, Q5 polymerase, and dNTPs at the recommended concentration. Primers that annealed directly up and downstream of the barcode and contained dual indexes to prevent “index swapping” on the Illumina HiSeq 4000 were added at 0.4 µM^95,96^. Primer sequences are listed in Table S3. To amplify barcodes, PCR mixes were heated to 98°C for 4 min followed by 25 cycles of 98°C for 30 s, 55°C for 30 s, and 72°C for 30 s. At the conclusion of 25 cycles, samples were incubated at 72°C for 5 min. PCR products were checked for the correct size on a 1% agarose gel, combined, and then purified using the DNA Clean & Concentrator-5 kit (Zymo Research). The combined, purified PCR products were quantified using the Qubit dsDNA HS assay on a Qubit 2.0 (Invitrogen) and visualized on an Agilent Fragment Analyzer at the Biotechnology Resource Center at Cornell University. Barcodes were sequenced on a NextSeq 500 (Illumina) with 75 bp single-end reads at the Cornell University Biotechnology Resource Center.

Fitness analysis was performed with the RBseq v. 1.1.9 pipeline (https://github.com/stcoradetti/RBseq)^26,52^ using the default settings comparing the barcode abundance in the final experimental populations to the barcode abundance in the paired starting population for each replicate. To avoid investigating pleiotropic mutants, we also computed fitness scores comparing the barcode abundance of samples exposed to VMM NH_4_Cl + 2% glucose supplemented with ammonium-phosphate buffer with 6.29 mM copper sulfate, 9 mM zinc sulfate, or 3.33 mM copper sulfate plus 3.19 mM zinc sulfate to the barcode abundance of samples exposed to VMM NH_4_Cl + 2% glucose supplemented with ammonium-phosphate buffer by running RBseq_Calculate_Fitness.py 1.1.9 (https://github.com/stcoradetti/RBseq)^26,52^ using the default settings with the paired VMM NH_4_Cl + 2% glucose supplemented with ammonium-phosphate buffer replicate in place of the T0. To identify copper plus zinc-specific mutants, we also used RBseq_Calculate_Fitness.py 1.1.9 (https://github.com/stcoradetti/RBseq)^26,52^ using the default settings to compute fitness scores comparing the barcode abundance of samples exposed to VMM NH_4_Cl + 2% glucose supplemented with ammonium-phosphate buffer and 3.33 mM copper sulfate plus 3.19 mM zinc sulfate to the barcode abundance of samples exposed to VMM NH_4_Cl + 2% glucose supplemented with ammonium-phosphate buffer and 6.29 mM copper sulfate or 9 mM zinc sulfate as the paired T0. To identify genes that cause metal-specific fitness phenotypes, we required that genes have a fitness of less than -2 or greater than 2 and p_adj_ < 0.05 relative to both the starting population and the buffer condition. Raw sequencing data are available in the NCBI Sequence Read Archive (SRA) (BioProject accession: PRJNA1536535).

## Statistical significance tests

Statistical analyses were performed in Rstudio (v2026.01.0+392; R v4.5.3), except for the BarSeq analysis which was run with Python (v 3.12.8). Statistical significance of all growth, metal accumulation, kill-curve, and RT-qPCR data was determined using either a one-way ANOVA with a Dunnett’s or Tukey’s post-hoc test, a two-way ANOVA with a Tukey’s post-hoc test, or a two-tailed two-sample *t*-test, as appropriate for the data being analyzed, and is indicated in the respective figure legend. Statistics on synergy calculations were performed using the SynergyFinder Plus R package v. 3.18.0^28^. Statistical significance for differential expression analyses was determined using DESeq2^89^. Functional enrichment analyses were performed using clusterprofiler^91^, as detailed above. Statistical significance for high-throughput fitness assays was determined using RBseq v. 1.1.9 (https://github.com/stcoradetti/RBseq)^26,52^. All experiments were performed with a minimum of three biological replicates.

## Data availability

Raw RNAseq data are available in GEO at the NCBI through accession numbers GSE348969 (*N. crassa*) and GSE348884 (*R. toruloides*). Processed RNAseq data is available in Datasets S1-S4. Raw barcode sequencing data are available on SRA at the NCBI through BioProject accession number PRJNA1536535. Processed BarSeq data are available in Dataset S5. The numerical values used to generate all other graphs are in Dataset S6. Strains constructed in this study are available upon request.

## ACKNOWLEDGEMENTS

This work was supported by a grant from VM Agritech to L.B.H. VM Agritech was not involved in the conceptualization, experimental design, data collection, data analysis, data interpretation, decision to publish, or preparation of the manuscript. J.D.K. was supported by a National Institutes of Health Ruth L. Kirschstein National Research Service Award Institutional Research Training Grant T32AI145821. This study is based on work supported by the National Science Foundation (NSF) Graduate Research Fellowship to R.E.S. under grant number DGE-2139899. A portion of this effort was supported by XLEAP at CHESS, NSF-BIO 2330043. This work utilized the Cornell Institute of Biotechnology Biotechnology Resource Center Genomics Facility (RRID:SCR_021727) and BioHPC (RRID:SCR_021757).

## REFERENCES

1. Denning, D. W. Global incidence and mortality of severe fungal disease. Lancet Infect. Dis. 24, e428–e438 (2024).

2. Savary, S. et al. The global burden of pathogens and pests on major food crops. *Nat*. Ecol. Evol. 3, 430–439 (2019).

3. van Rhijn, N. & Rhodes, J. Evolution of antifungal resistance in the environment. Nat. Microbiol. 10, 1804–1815 (2025).

4. Perlin, D. S., Rautemaa-Richardson, R. & Alastruey-Izquierdo, A. The global problem of antifungal resistance: prevalence, mechanisms, and management. Lancet Infect. Dis. 17, e383–e392 (2017).

5. Fisher, M. C. et al. A one health roadmap towards understanding and mitigating emerging Fungal Antimicrobial Resistance: fAMR. Npj Antimicrob. Resist. 2, 36 (2024).

6. Johnson, G. F. The Early History of Copper Fungicides. Agric. Hist. 9, 67–79 (1935).

7. La Torre, A., Iovino, V. & Caradonia, F. Copper in plant protection: current situation and prospects. Phytopathol. Mediterr. 57, 201–236 (2018).

8. Solioz, M. Copper Toxicity. in Copper and Bacteria: Evolution, Homeostasis and Toxicity (ed. Solioz, M.) 11–19 (Springer International Publishing, Cham, 2018). doi:10.1007/978-3-319-94439-5_2.

9. Zuily, L. et al. Copper Induces Protein Aggregation, a Toxic Process Compensated by Molecular Chaperones. mBio 13, e03251–21 (2022).

10. Smith, A. D., Logeman, B. L. & Thiele, D. J. Copper Acquisition and Utilization in Fungi. Annu. Rev. Microbiol. 71, 597–623 (2017).

11. Wiebelhaus, N., Zaengle-Barone, J. M., Hwang, K. K., Franz, K. J. & Fitzgerald, M. C. Protein Folding Stability Changes Across the Proteome Reveal Targets of Cu Toxicity in E. coli. ACS Chem. Biol. 16, 214–224 (2021).

12. Saporito-Magriñá, C. M. et al. Copper-induced cell death and the protective role of glutathione: the implication of impaired protein folding rather than oxidative stress†. Metallomics 10, 1743–1754 (2018).

13. Macomber, L. & Imlay, J. A. The iron-sulfur clusters of dehydratases are primary intracellular targets of copper toxicity. Proc. Natl. Acad. Sci. 106, 8344–8349 (2009).

14. Gunther, M. R., Hanna, P. M., Mason, R. P. & Cohen, M. S. Hydroxyl Radical Formation from Cuprous Ion and Hydrogen Peroxide: A Spin-Trapping Study. Arch. Biochem. Biophys. 316, 515–522 (1995).

15. Neaman, A. et al. Copper contamination in agricultural soils: A review of the effects of climate, soil properties, and prolonged copper pesticide application in vineyards and orchards. Plant Soil Environ. 70, 407–417 (2024).

16. Chandrangsu, P., Rensing, C. & Helmann, J. D. Metal homeostasis and resistance in bacteria. Nat. Rev. Microbiol. 15, 338–350 (2017).

17. McDevitt, C. A. et al. A Molecular Mechanism for Bacterial Susceptibility to Zinc. PLOS Pathog. 7, e1002357 (2011).

18. Xu, Z. et al. Zinc excess increases cellular demand for iron and decreases tolerance to copper in *Escherichia coli*. J. Biol. Chem. 294, 16978–16991 (2019).

19. Song, W. et al. Role of the dissolved zinc ion and reactive oxygen species in cytotoxicity of ZnO nanoparticles. Toxicol. Lett. 199, 389–397 (2010).

20. Shen, C. et al. Relating Cytotoxicity, Zinc Ions, and Reactive Oxygen in ZnO Nanoparticle–Exposed Human Immune Cells. Toxicol. Sci. 136, 120–130 (2013).

21. Sadhasivam, S. et al. Antifungal effect and mode of activity of zinc chloride against toxigenic fungus *Aspergillus flavus*. Food Res. Int. 208, 116160 (2025).

22. Naranjo, E. et al. Zinkicide Is a ZnO-Based Nanoformulation with Bactericidal Activity against Liberibacter crescens in Batch Cultures and in Microfluidic Chambers Simulating Plant Vascular Systems. Appl. Environ. Microbiol. 86, e00788–20 (2020).

23. Sun, Q., Li, J. & Le, T. Zinc Oxide Nanoparticle as a Novel Class of Antifungal Agents: Current Advances and Future Perspectives. J. Agric. Food Chem. 66, 11209–11220 (2018).

24. Hamzah Saleem, M., Usman, K., Rizwan, M., Al Jabri, H. & Alsafran, M. Functions and strategies for enhancing zinc availability in plants for sustainable agriculture. Front. Plant Sci. 13, (2022).

25. Nielsen, F. H. History of Zinc in Agriculture. Adv. Nutr. 3, 783–789 (2012).

26. Coradetti, S. T. et al. Functional genomics of lipid metabolism in the oleaginous yeast Rhodosporidium toruloides. eLife 7, e32110 (2018).

27. Vogel, H. J. A convenient growth medium for Neurospora (medium N). Microb. Genet Bull 13, 42–43 (1956).

28. Zheng, S. et al. SynergyFinder Plus: Toward Better Interpretation and Annotation of Drug Combination Screening Datasets. Genomics Proteomics Bioinformatics 20, 587–596 (2022).

29. Yadav, B., Wennerberg, K., Aittokallio, T. & Tang, J. Searching for Drug Synergy in Complex Dose–Response Landscapes Using an Interaction Potency Model. Comput. Struct. Biotechnol. J. 13, 504–513 (2015).

30. Douglas, L. M. & Konopka, J. B. Plasma membrane architecture protects Candida albicans from killing by copper. PLOS Genet. 15, e1007911 (2019).

31. Quaranta, D. et al. Mechanisms of Contact-Mediated Killing of Yeast Cells on Dry Metallic Copper Surfaces. Appl. Environ. Microbiol. 77, 416–426 (2011).

32. Hwang, Y.-J., Kim, Y.-Y., Chang, J.-Y. & Kho, H.-S. Candidacidal activities of zinc compounds based on incubation time, pH, and ionic strength – An in vitro study. J. Dent. Sci. 20, 885–894 (2025).

33. Djearamane, S., et al. Antifungal Properties of Zinc Oxide Nanoparticles on Candida albicans. Coatings 12, (2022).

34. The Gene Ontology Consortium. The Gene Ontology knowledgebase in 2026. Nucleic Acids Res. 54, D1779–D1792 (2026).

35. Ashburner, M. et al. Gene Ontology: tool for the unification of biology. Nat. Genet. 25, 25–29 (2000).

36. Kanehisa, M. & Goto, S. KEGG: Kyoto Encyclopedia of Genes and Genomes. Nucleic Acids Res. 28, 27–30 (2000).

37. Kanehisa, M. Toward understanding the origin and evolution of cellular organisms. Protein Sci. 28, 1947–1951 (2019).

38. Kanehisa, M., Furumichi, M., Sato, Y., Matsuura, Y. & Ishiguro-Watanabe, M. KEGG: biological systems database as a model of the real world. Nucleic Acids Res. 53, D672–D677 (2025).

39. Franklin, C. C. et al. Structure, function, and post-translational regulation of the catalytic and modifier subunits of glutamate cysteine ligase. Mol. Aspects Med. 30, 86–98 (2009).

40. Hu, C. et al. Heat shock proteins: Biological functions, pathological roles, and therapeutic opportunities. MedComm 3, e161 (2022).

41. Huh, W.-K. et al. Global analysis of protein localization in budding yeast. Nature 425, 686–691 (2003).

42. Breker, M., Gymrek, M. & Schuldiner, M. A novel single-cell screening platform reveals proteome plasticity during yeast stress responses. J. Cell Biol. 200, 839–850 (2013).

43. Breker, M., Gymrek, M., Moldavski, O. & Schuldiner, M. LoQAtE—Localization and Quantitation ATlas of the yeast proteomE. A new tool for multiparametric dissection of single-protein behavior in response to biological perturbations in yeast. Nucleic Acids Res. 42, D726–D730 (2014).

44. Dubreuil, B. et al. YeastRGB: comparing the abundance and localization of yeast proteins across cells and libraries. Nucleic Acids Res. 47, D1245–D1249 (2019).

45. Shi, Y. et al. The dynamic roles of intracellular vacuoles in heavy metal detoxification by Rhodotorula mucilaginosa. J. Appl. Microbiol. 135, lxae241 (2024).

46. Szczypka, M. S., Zhu, Z., Silar, P. & Thiele, D. J. Saccharomyces cerevisiae mutants altered in vacuole function are defective in copper detoxification and iron-responsive gene transcription. Yeast 13, 1423–1435 (1997).

47. Cho, M. et al. Vacuolar zinc transporter Zrc1 is required for detoxification of excess intracellular zinc in the human fungal pathogen Cryptococcus neoformans. J. Microbiol. 56, 65–71 (2018).

48. Mendoza-Cózatl, D. G. et al. Tonoplast-localized Abc2 Transporter Mediates Phytochelatin Accumulation in Vacuoles and Confers Cadmium Tolerance *. J. Biol. Chem. 285, 40416–40426 (2010).

49. Price, M. N. et al. Indirect and suboptimal control of gene expression is widespread in bacteria. Mol. Syst. Biol. 9, MSB201316 (2013).

50. Protzko, R. J. et al. Genomewide and Enzymatic Analysis Reveals Efficient d-Galacturonic Acid Metabolism in the Basidiomycete Yeast Rhodosporidium toruloides. mSystems 4, 10.1128/msystems.00389-19 (2019).

51. Giaever, G. et al. Functional profiling of the Saccharomyces cerevisiae genome. Nature 418, 387–391 (2002).

52. Kim, J. et al. Multi-Omics Driven Metabolic Network Reconstruction and Analysis of Lignocellulosic Carbon Utilization in Rhodosporidium toruloides. Front. Bioeng. Biotechnol. 8, (2021).

53. Jensen, P. A., Zhu, Z. & Opijnen, T. van. Antibiotics Disrupt Coordination between Transcriptional and Phenotypic Stress Responses in Pathogenic Bacteria. Cell Rep. 20, 1705–1716 (2017).

54. Huberman, L. B., et al. Barcoded mutant library enables high-throughput functional genomics in a filamentous fungus. Proc. Natl. Acad. Sci. 123, e2616888123 (2026).

55. Madeira, F. et al. The EMBL-EBI Job Dispatcher sequence analysis tools framework in 2024. 10.1093/nar/gkae241.

56. Stoj, C. & Kosman, D. J. Cuprous oxidase activity of yeast Fet3p and human ceruloplasmin: implication for function. FEBS Lett. 554, 422–426 (2003).

57. Askwith, C. et al. The FET3 gene of S. cerevisiae encodes a multicopper oxidase required for ferrous iron uptake. Cell 76, 403–410 (1994).

58. Felice, M. R. et al. Post-transcriptional Regulation of the Yeast High Affinity Iron Transport System *. J. Biol. Chem. 280, 22181–22190 (2005).

59. Shi, X., Stoj, C., Romeo, A., Kosman, D. J. & Zhu, Z. Fre1p Cu2+ Reduction and Fet3p Cu1+ Oxidation Modulate Copper Toxicity in *Saccharomyces cerevisiae*\*. J. Biol. Chem. 278, 50309–50315 (2003).

60. Gross, C., Kelleher, M., Iyer, V. R., Brown, P. O. & Winge, D. R. Identification of the Copper Regulon in *Saccharomyces cerevisiae* by DNA Microarrays*. J. Biol. Chem. 275, 32310–32316 (2000).

61. Gaspar-Cordeiro, A., Marques Caetano, S., Amaral, C., Rodrigues-Pousada, C. & Pimentel, C. Ace1 prevents intracellular copper accumulation by regulating Fet3 expression and thereby restricting Aft1 activity. FEBS J. 285, 1861–1872 (2018).

62. Ruehle, G. D. Developments In Tomato Production In Florida: The Use Of Zinc Salts With Copper Fungicides On Tomatoes In Dade County. Proc. Fla. State Hortic. Soc. 55, 133–133 (1942).

63. Tsiridis, V. et al. Interactive toxic effects of heavy metals and humic acids on *Vibrio fischeri*. Ecotoxicol. Environ. Saf. 63, 158–167 (2006).

64. Cordeiro, L. M. et al. Toxicity of copper and zinc alone and in combination in *Caenorhabditis elegans* model of Huntington’s disease and protective effects of rutin. NeuroToxicology 97, 120–132 (2023).

65. Fagnano, M. et al. Copper accumulation in agricultural soils: Risks for the food chain and soil microbial populations. Sci. Total Environ. 734, 139434 (2020).

66. Department of Agriculture, A. M. S. The National List of Allowed and Prohibited Substances. C.F.R. vol. 7 (2000).

67. Deng, W. et al. Biological remediation mechanism and applications of *Rhodotorula mucilaginosa* for heavy metal pollution. *Environ*. Technol. Innov. 38, 104179 (2025).

68. Cavelius, P. et al. Adaptation of Proteome and Metabolism in Different Haplotypes of Rhodosporidium toruloides during Cu(I) and Cu(II) Stress. Microorganisms 11, (2023).

69. Kan, G. et al. Copper stress response in yeast Rhodotorula mucilaginosa AN5 isolated from sea ice, Antarctic. MicrobiologyOpen 8, e00657 (2019).

70. Khemiri, I., Tebbji, F. & Sellam, A. Transcriptome Analysis Uncovers a Link Between Copper Metabolism, and Both Fungal Fitness and Antifungal Sensitivity in the Opportunistic Yeast Candida albicans. Front. Microbiol. 11, (2020).

71. Sun, T. et al. Proteomic Analysis of Copper Toxicity in Human Fungal Pathogen Cryptococcus neoformans. Front. Cell. Infect. Microbiol. 11, (2021).

72. Cain, A. K. et al. A decade of advances in transposon-insertion sequencing. Nat. Rev. Genet. 21, 526–540 (2020).

73. Torres, M., Paszti, S. & Eberl, L. Shedding light on bacteria–host interactions with the aid of TnSeq approaches. mBio 15, e00390–24 (2024).

74. Colot, H. V. et al. A high-throughput gene knockout procedure for Neurospora reveals functions for multiple transcription factors. Proc. Natl. Acad. Sci. 103, 10352–10357 (2006).

75. McCluskey, K., Wiest, A. & Plamann, M. The Fungal Genetics Stock Center: a repository for 50 years of fungal genetics research. J. Biosci. 35, 119–126 (2010).

76. Zhang, S., Ito, M., Skerker, J. M., Arkin, A. P. & Rao, C. V. Metabolic engineering of the oleaginous yeast Rhodosporidium toruloides IFO0880 for lipid overproduction during high-density fermentation. Appl. Microbiol. Biotechnol. 100, 9393–9405 (2016).

77. Abbott, E. P., Ianiri, G., Castoria, R. & Idnurm, A. Overcoming recalcitrant transformation and gene manipulation in Pucciniomycotina yeasts. Appl. Microbiol. Biotechnol. 97, 283–295 (2013).

78. Reyes-Chavez, B., Kerkaert, J. D. & Huberman, L. B. The carbohydrate utilization regulator Cbr1 coordinates nutrient-specific gene activation with selective carbon catabolite repression in a basidiomycete yeast. PLOS Biol. 24, e3003983 (2026).

79. Coradetti, S. T. et al. Engineering transcriptional regulation of pentose metabolism in Rhodosporidium toruloides for improved conversion of xylose to bioproducts. Microb. Cell Factories 22, 144 (2023).

80. Wang, Y. et al. Cloning and evaluation of different constitutive promoters in the oleaginous yeast Rhodosporidium toruloides. Yeast 33, 99–106 (2016).

81. Solé, V. A., Papillon, E., Cotte, M., Walter, Ph. & Susini, J. A multiplatform code for the analysis of energy-dispersive X-ray fluorescence spectra. Spectrochim. Acta Part B At. Spectrosc. 62, 63–68 (2007).

82. Wu, V. W. et al. The regulatory and transcriptional landscape associated with carbon utilization in a filamentous fungus. Proc. Natl. Acad. Sci. 117, 6003–6013 (2020).

83. Chen, S., Zhou, Y., Chen, Y. & Gu, J. fastp: an ultra-fast all-in-one FASTQ preprocessor. Bioinformatics 34, i884–i890 (2018).

84. Galagan, J. E. et al. The genome sequence of the filamentous fungus Neurospora crassa. Nature 422, 859–868 (2003).

85. Dobin, A. et al. STAR: ultrafast universal RNA-seq aligner. Bioinformatics 29, 15–21 (2013).

86. Danecek, P. et al. Twelve years of SAMtools and BCFtools. GigaScience 10, giab008 (2021).

87. Tange, O. GNU Parallel: The Command-Line Power Tool. USENIX Mag. 42–47 (2011).

88. Putri, G. H., Anders, S., Pyl, P. T., Pimanda, J. E. & Zanini, F. Analysing high-throughput sequencing data in Python with HTSeq 2.0. Bioinformatics 38, 2943–2945 (2022).

89. Love, M. I., Huber, W. & Anders, S. Moderated estimation of fold change and dispersion for RNA-seq data with DESeq2. Genome Biol. 15, 550 (2014).

90. Gu, Z. Complex heatmap visualization. iMeta 1, e43 (2022).

91. Xu, S. et al. Using clusterProfiler to characterize multiomics data. Nat. Protoc. 19, 3292–3320 (2024).

92. The UniProt Consortium. UniProt: the Universal Protein Knowledgebase in 2025. Nucleic Acids Res. 53, D609–D617 (2025).

93. Basenko, E. Y. et al. What is new in FungiDB: a web-based bioinformatics platform for omics-scale data analysis for fungal and oomycete species. Genetics 227, iyae035 (2024).

94. Alvarez-Jarreta, J. et al. VEuPathDB: the eukaryotic pathogen, vector and host bioinformatics resource center in 2023. Nucleic Acids Res. 52, D808–D816 (2024).

95. Price, M. N., et al. Oxidative Pathways of Deoxyribose and Deoxyribonate Catabolism. mSystems 4, 10.1128/msystems.00297-18 (2019).

96. Costello, M. et al. Characterization and remediation of sample index swaps by non-redundant dual indexing on massively parallel sequencing platforms. BMC Genomics 19, 332 (2018).

