## Supplemental Appendix for "High-throughput functional genomics and transcriptomics reveal molecular mechanisms regulating the synergistic fungicidal activity of copper and zinc"

### SI FIGURES

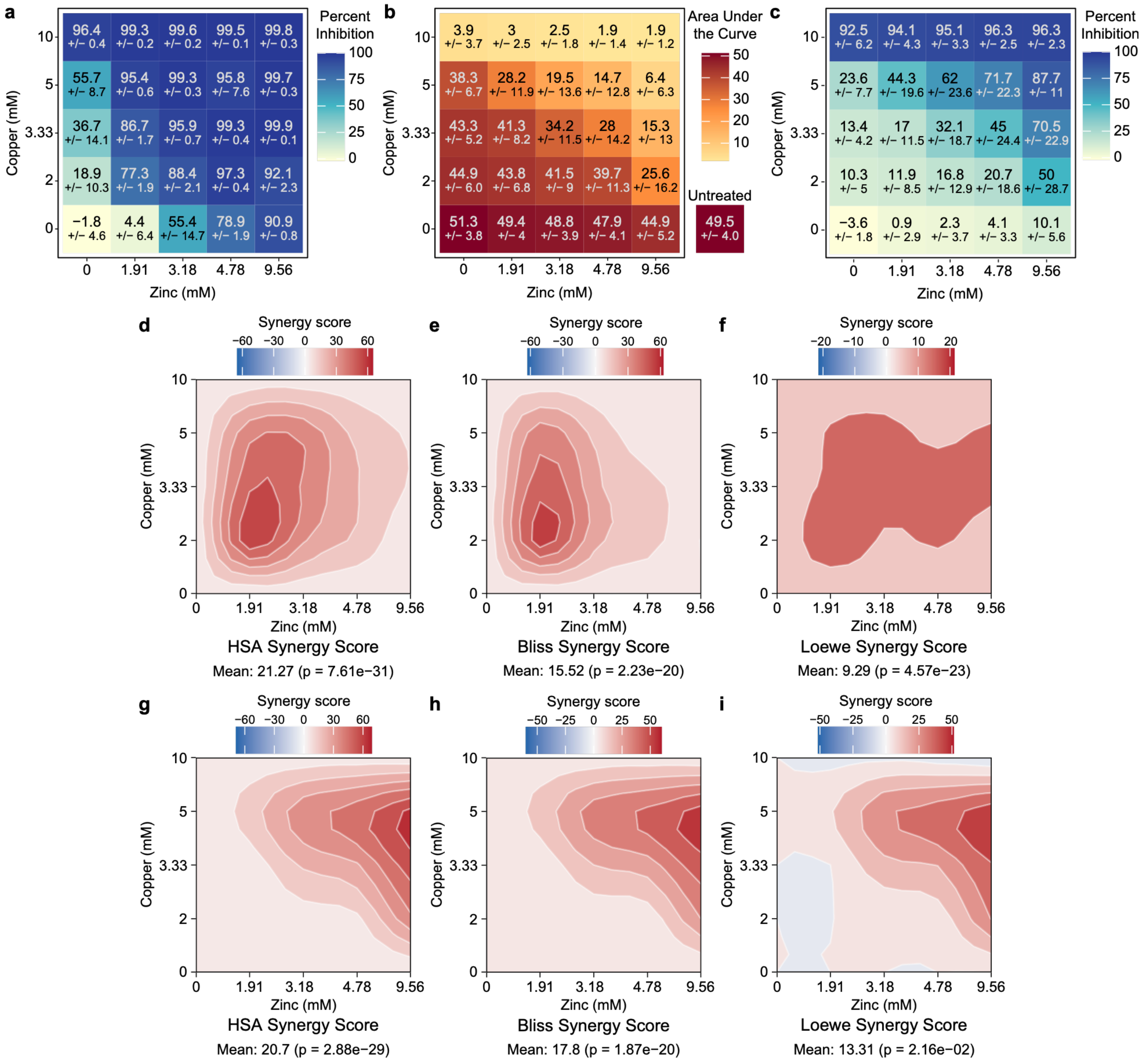


**Fig S1. Copper and zinc act synergistically to reduce fungal growth. (A)** Percent growth inhibition of *N. crassa* treated with the indicated concentrations of copper sulfate and zinc sulfate in an ammonium-phosphate buffer relative to growth in media without any added ammonium-phosphate buffer, copper sulfate, or zinc sulfate. **(B)** Area under the curve calculations, from growth curves in Fig 1B, of *R. toruloides* treated with the indicated concentrations of copper sulfate and zinc sulfate in an ammonium-phosphate buffer. **(C)** Percent growth inhibition of *R. toruloides* treated with the indicated concentrations of copper sulfate and zinc sulfate in an ammonium-phosphate buffer relative to growth on media without any added ammonium-phosphate buffer, copper sulfate, or zinc sulfate. **(D-I)** (**D** and **G**) Highest single agent (HSA), (**E** and **H**) Bliss, and (**F** and **I**) Loewe synergy plots for the percent inhibition of *N. crassa* (**E-F**) and *R. toruloides* (**G-I**) by the indicated concentration of copper sulfate and zinc sulfate. To generate percent inhibition values, the *N. crassa* biomass data (from Fig 1A) and *R. toruloides* area under the curve values (from Fig S1B) were normalized to a media-only control for each experiment. For (**A-C**) the top values are the mean percent inhibition (**A** and **C**) or area under the curve (**B**), and the lower numbers are the standard deviation of (**A**) four or (**B-C**) five (single metal treatments) or six biological replicates (all other groups). For (**D-I**) the average synergy score and p-value are displayed. Synergy calculations and contour plot generation was performed using the SynergyFinder+ R package[1].

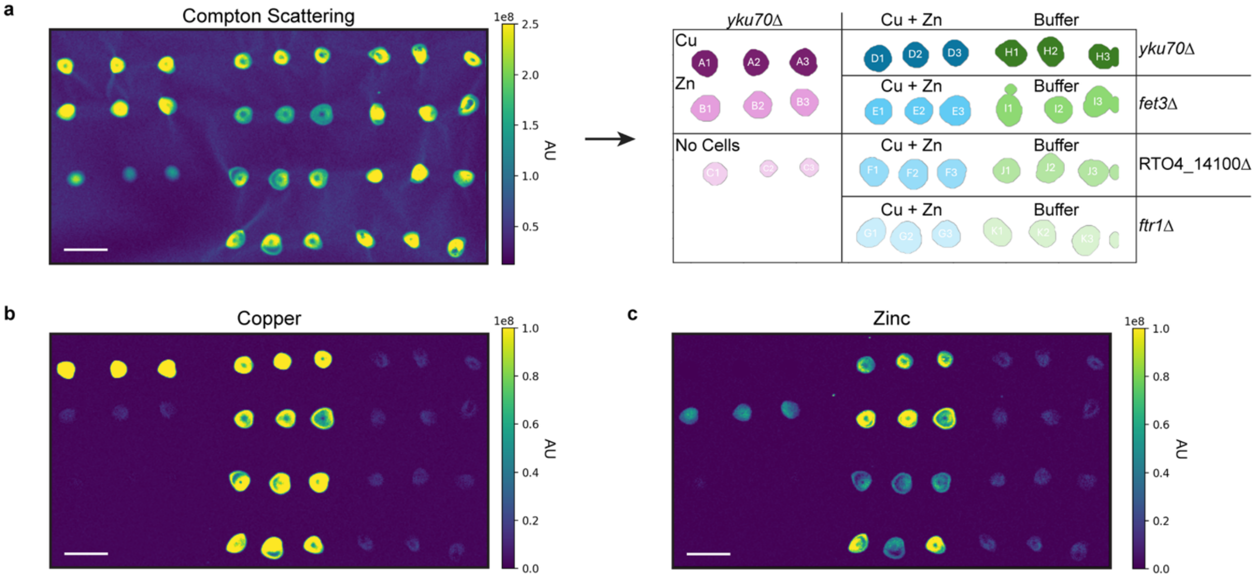


**Fig S2. Quantification of copper and zinc using x-ray fluorescence imaging. (A)** Compton scattering heatmap (left) was used to generate a binary mask (right) for quantification of total fluorescence intensity within each indicated region to generate bar graphs in Fig 2 and Fig 6F-6G. The binary mask is labeled with the corresponding strain and treatment for each spot of cells or media: A1-A3 – *yku70*Δ cells exposed to 5 mM copper sulfate in ammonium-phosphate buffer; B1-B3 – *yku70*Δ cells exposed to 4.78 mM zinc sulfate in ammonium-phosphate buffer; C1-C3 – Media + ammonium-phosphate buffer (no cells); D1-D3 – *yku70*Δ cells exposed to 5 mM copper sulfate + 4.78 mM zinc sulfate in ammonium-phosphate buffer; E1-E3 – *fet3*Δ cells exposed to 5 mM copper sulfate + 4.78 mM zinc sulfate in ammonium-phosphate buffer; F1-F3 –RTO4_14100Δ cells exposed to 5 mM copper sulfate + 4.78 mM zinc sulfate in ammonium-phosphate buffer; G1-G3 – *ftr1*Δ cells exposed to 5 mM copper sulfate + 4.78 mM zinc sulfate in ammonium-phosphate buffer; H1-H3 – *yku70*Δ cells exposed to ammonium-phosphate buffer; I1-I3 – *fet3*Δ cells exposed to ammonium-phosphate buffer; J1-J3 – RTO4_14100Δ cells exposed to ammonium-phosphate buffer; K1-K3 – *ftr1*Δ cells exposed to ammonium-phosphate buffer. Each spot contains an equal amount of washed cells dried onto Kapton film for quantification of cellular metal concentration using x-ray fluorescence imaging and corresponds to a single biological replicate. Spots containing an equal volume of media supplemented with ammonium-phosphate buffer were included as controls for levels of copper and zinc present in the media (spots C1-C3). **(B-C)** Quantification heatmaps of (**B**) copper and (**C**) zinc concentration. Scale bar is 1 cm. AU stands for arbitrary units.

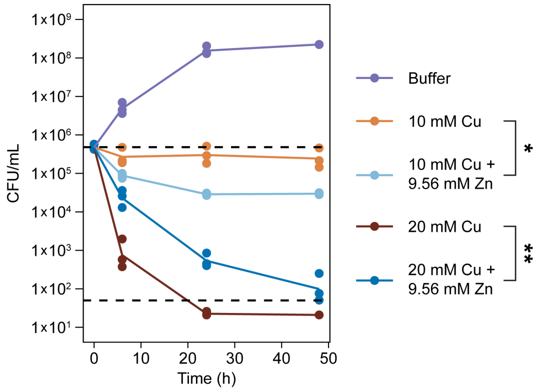


**Fig S3. Copper and zinc in combination is fungicidal.** Time-kill curve of *R. toruloides* treated with the indicated concentrations of copper sulfate and/or copper sulfate plus zinc sulfate in an ammonium-phosphate buffer or the ammonium-phosphate buffer alone. Colony forming units (CFU) per mL were quantified after 0 h, 6 h, 24 h, and 48 h of treatment. Upper dashed line indicates the average starting CFU/mL ($4.8x10^5$ CFU/mL). Lower dashed line indicates the limit of detection (50 CFU/mL). All samples below the limit of detection were assigned a value of 20 for the purposes of visualization on a log scale. *$p < 0.05$ and **$p < 10^{-8}$ as determined by pairwise comparison of estimated marginal means with a Tukey multiple comparison correction performed on $\log_{10}$(CFU/mL + 1) transformed data.

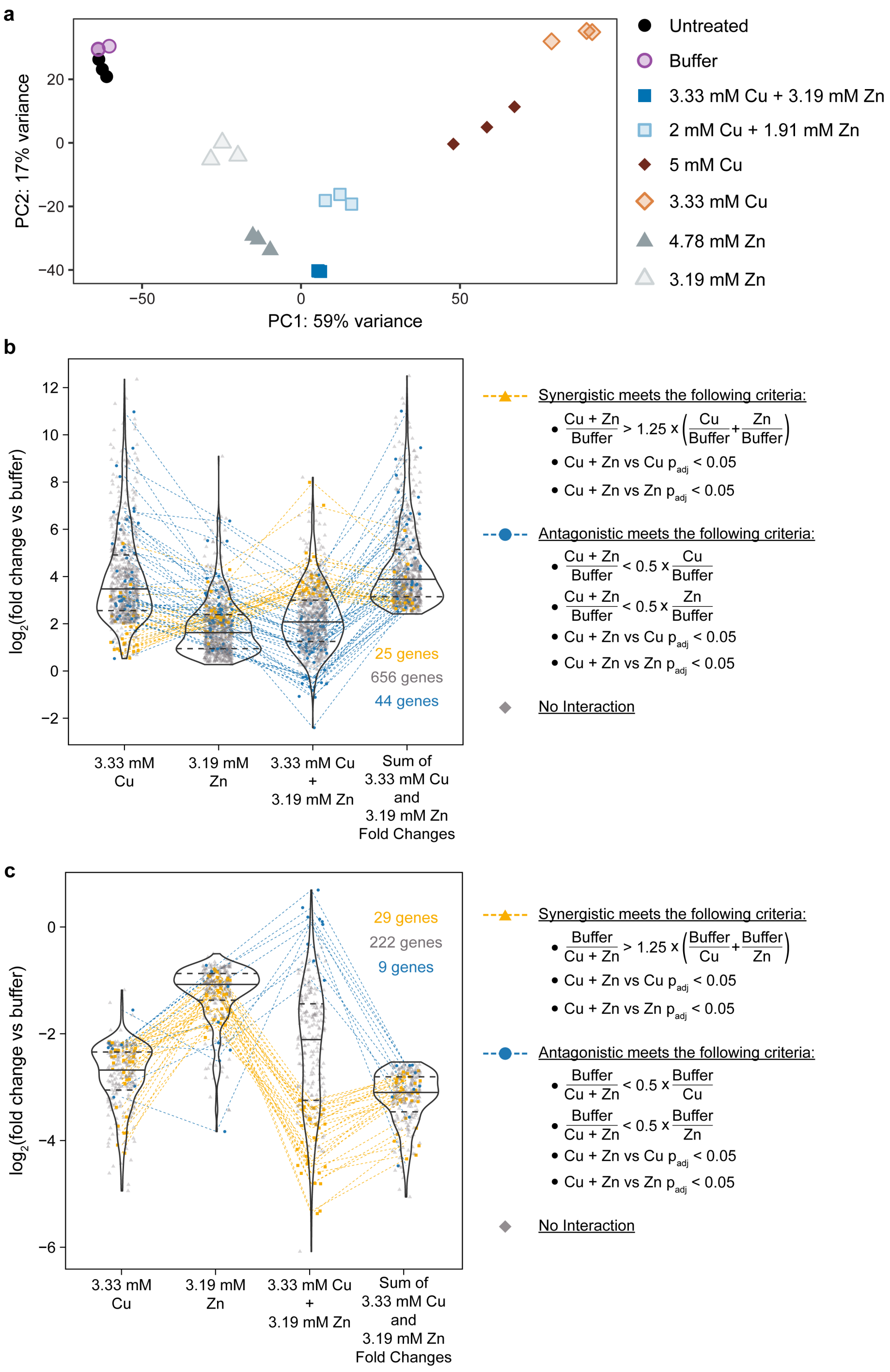


**Fig S4. Transcriptional profile of *N. crassa* treated with copper and zinc is more similar to treatment with zinc than copper. (A)** Principal component analysis of the variance stabilization transformed expression of the 2000 genes with the highest variance across all conditions in the *N. crassa* RNAseq dataset following a $\log_2$ transformation. **(B-C)** Analysis of genes significantly (**B**) increased or (**C**) decreased in both 3.33 mM copper sulfate and 3.19 mM zinc sulfate relative to treatment with buffer ($p_{adj} < 0.05$ for both conditions, with a four-fold or greater change in at least one of the two conditions). For genes increased in both conditions (**B**), the transcriptional response of a gene to 3.33 mM copper sulfate + 3.19 mM zinc sulfate (third group from left) was defined as synergistic (gold) if the magnitude of the fold change in its expression relative to buffer was at least 25% greater than the summed magnitude of the fold change in expression of that gene during exposure to 3.33 mM copper and 3.19 mM zinc individually (rightmost group) and the transcriptional response to 3.33 mM copper sulfate + 3.19 mM zinc sulfate was statistically significantly different ($p_{adj} < 0.05$) from both individual treatments. The transcriptional response of a gene to 3.33 mM copper sulfate + 3.19 mM zinc sulfate was defined as antagonistic (blue) if the magnitude of the fold change in its expression relative to buffer was less than half the magnitude of the fold change in expression during exposure to 3.33 mM copper and 3.19 mM zinc, individually, relative to buffer and the transcriptional response to 3.33 mM copper sulfate + 3.19 mM zinc sulfate was statistically significantly different ($p_{adj} < 0.05$) from both individual treatments. For genes decreased in both conditions (**C**), the inverse criteria were used for defining synergistic and antagonistic genes. All genes that did not meet these thresholds were defined to have no interaction (grey). The number of genes in each category is indicated in the corresponding text color. Statistical significance and fold changes were calculated using DESeq2[2].

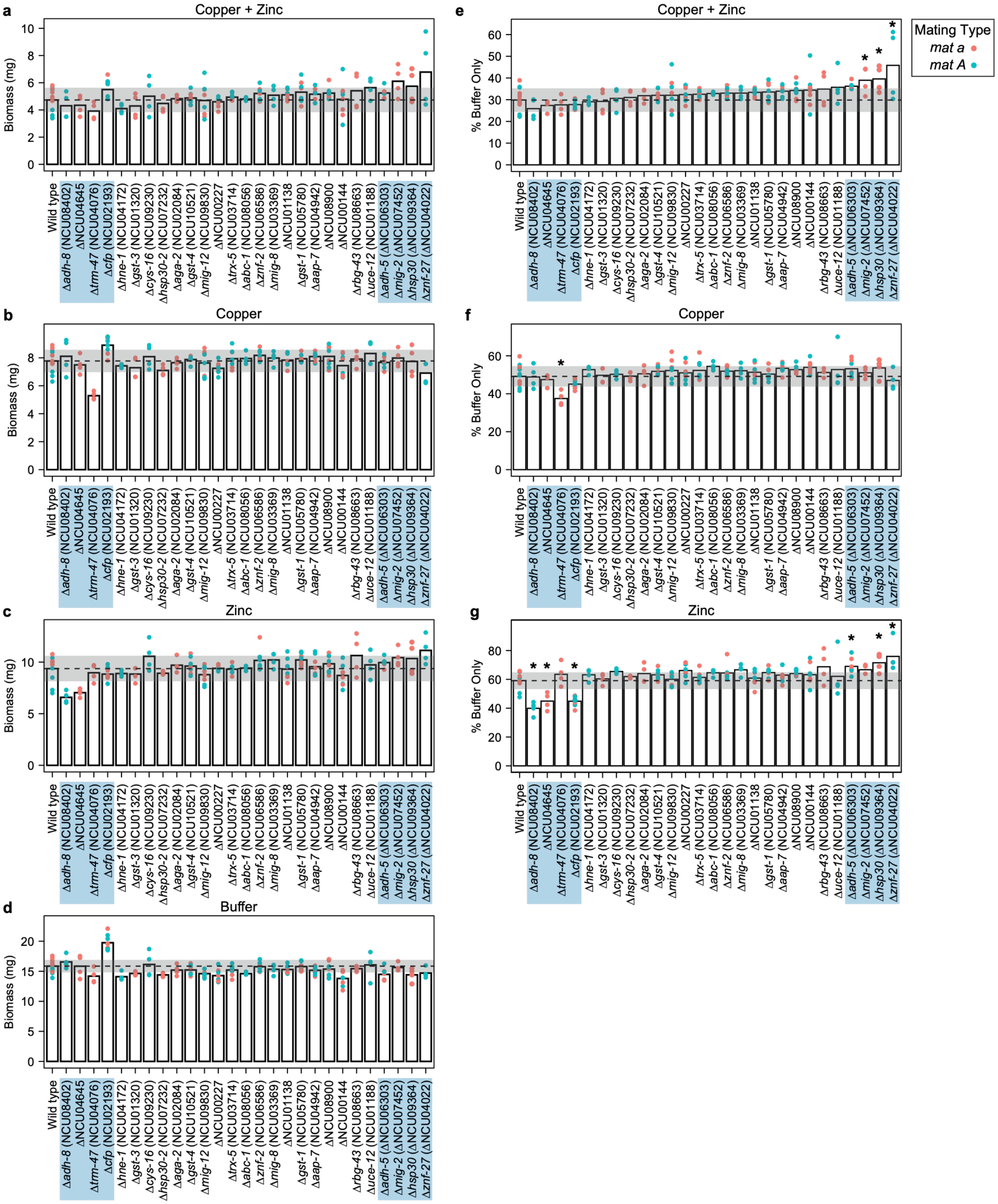


**Fig S5. Identification of *N. crassa* genes involved in resistance to copper and zinc in 3 mL cultures. (A-D)** Mycelial dry weight of *N. crassa* wild type cells and indicated deletion mutants directly inoculated into 3 mL of media supplemented with (**A**) 2 mM copper sulfate plus 1.91 mM zinc sulfate and ammonium-phosphate buffer, (**B**) 3.33 mM copper sulfate and ammonium-phosphate buffer, (**C**) 3.19 mM zinc sulfate and ammonium-phosphate buffer, and (**D**) ammonium-phosphate buffer and grown for two days. **(E-G)** Relative biomass accumulation of the indicated strains inoculated into media supplemented with ammonium-phosphate buffer with (**E**) 2 mM copper sulfate plus 1.91 mM zinc sulfate, (**F**) 3.33 mM copper sulfate, or (**G**) 3.19 mM zinc sulfate normalized to the biomass of the indicated strain inoculated into media supplemented with ammonium-phosphate buffer. Bars are averages and dots are individual data points colored by the mating type of the strain (red dots indicate *mat a* strains and blue dots indicate *mat A* strains) of at least four biological replicates or at least four biological replicates per mating type for genes in which the deletion mutant exists in both mating types in the *N. crassa* deletion collection[3,4]. No significant effect of mating type was observed. Dotted black lines indicate the average biomass accumulation or average normalized biomass accumulation and grey bands indicate standard deviations of 12 biological replicates (six biological replicates per mating type) of wild type cells exposed to the indicated condition. Deletion strains highlighted in blue were assessed for copper and zinc sensitivities in the secondary screen with 100 mL culture volumes (Fig 3D-F and S6). *$p < 0.05$ as determined by one-way ANOVA with a Dunnett's post-hoc test.

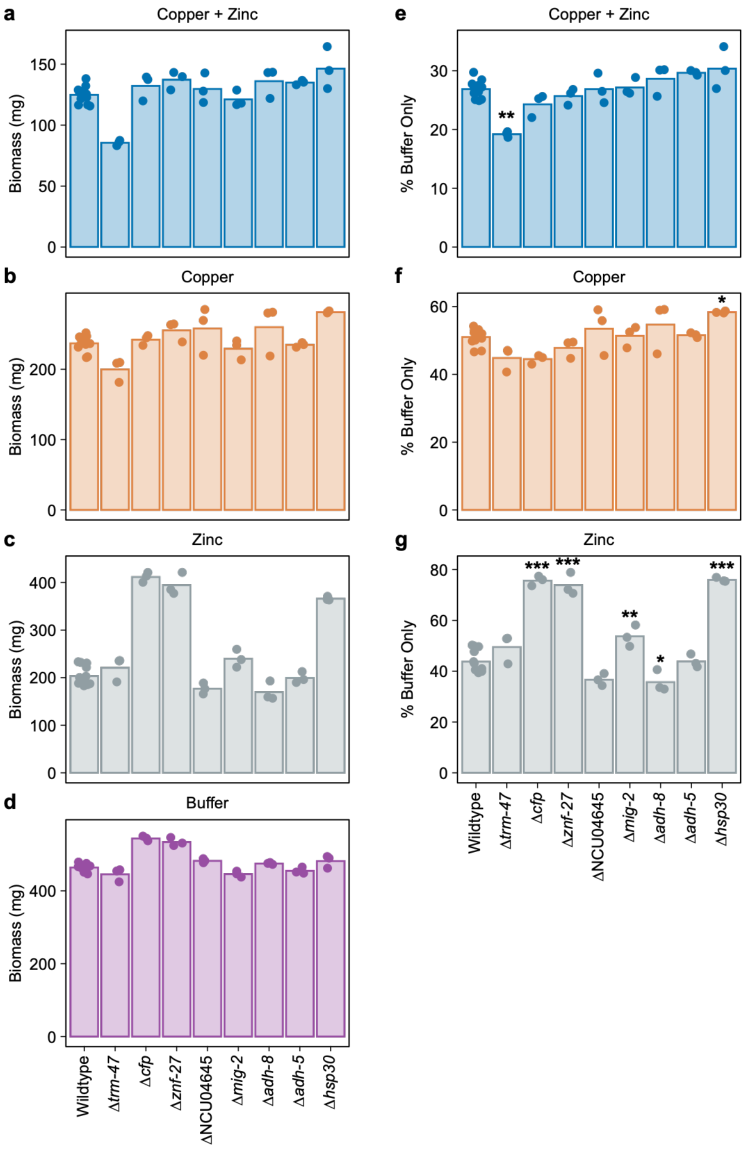


**Fig S6. Identification of *N. crassa* genes involved in resistance to copper and zinc in 100 mL cultures. (A-D)** Mycelial dry weight of *N. crassa* wild type cells and indicated deletion mutants inoculated into 100 mL of media supplemented with (**A**) 2 mM copper sulfate plus 1.91 mM zinc sulfate and ammonium-phosphate buffer, (**B**) 3.33 mM copper sulfate and ammonium-phosphate buffer, (**C**) 3.19 mM zinc sulfate and ammonium phosphate buffer, and (**D**) ammonium-phosphate buffer and grown for two days. **(E-G)** Relative biomass accumulation of the indicated strains inoculated into media supplemented with ammonium-phosphate buffer with (**E**) 2 mM copper sulfate plus 1.91 mM zinc sulfate, (**F**) 3.33 mM copper sulfate, or (**G**) 3.19 mM zinc sulfate normalized to the biomass of the indicated strain inoculated into media supplemented with ammonium-phosphate buffer. Bars are averages and dots are individual data points of at least three biological replicates. $^{*}p < 0.05$, $^{**}p < 10^{-5}$, and $^{***}p < 10^{-12}$ as determined by one-way ANOVA with a Dunnett's post-hoc test compared to wild type cells.

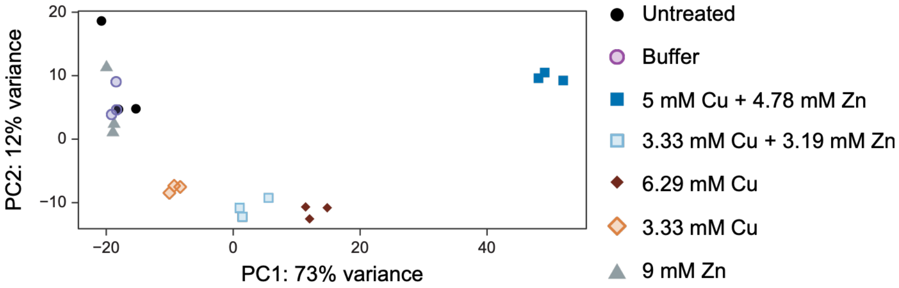


**Fig S7. Transcriptional profile of *R. toruloides* treated with copper and zinc is more similar to copper treatment than zinc treatment**. Principle components analysis of the variance stabilization transformed expression of the 1000 genes with the highest variance across all conditions in the *R. toruloides* RNAseq dataset following a $\log_2$ transformation.

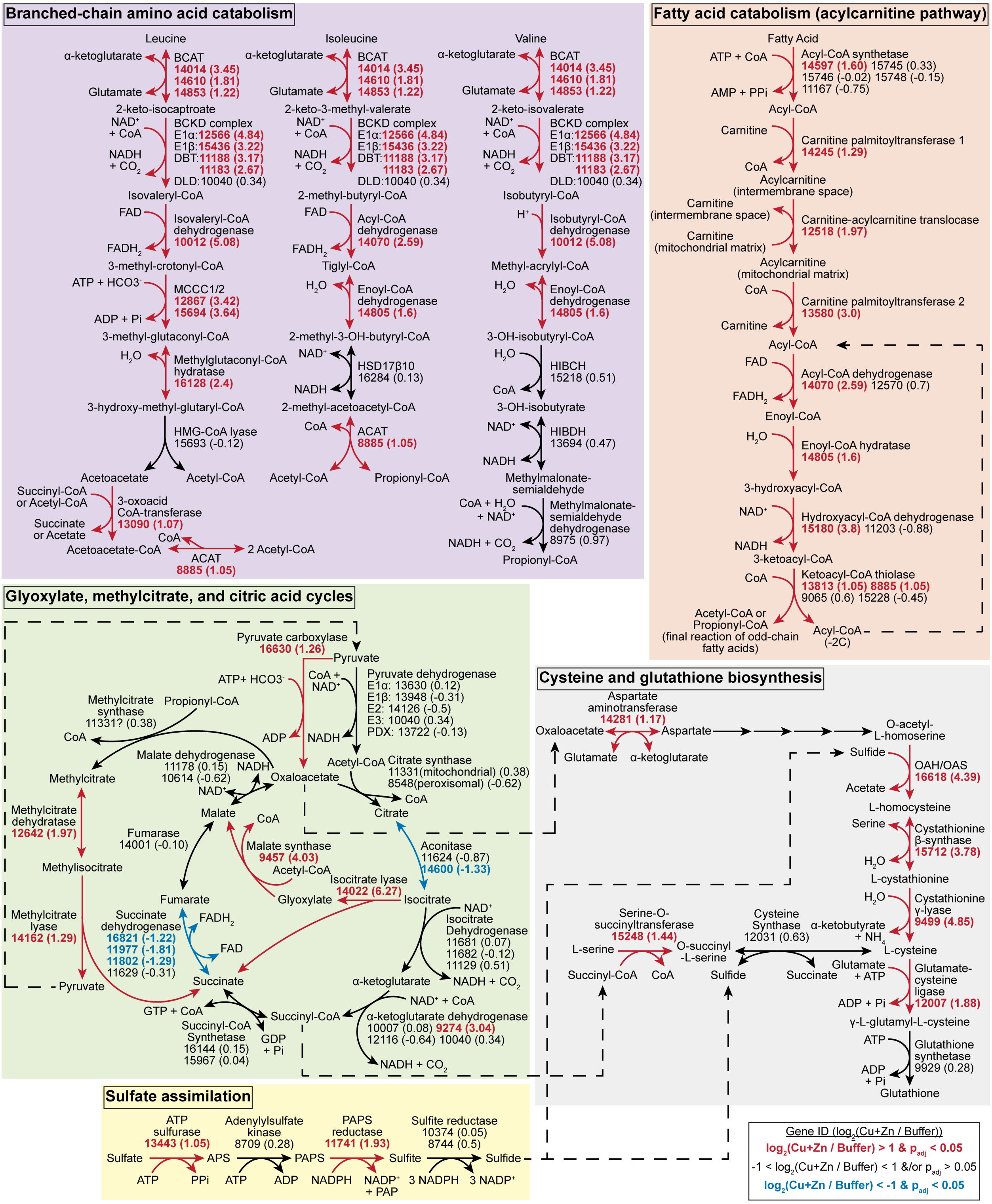


**Fig S8. In response to treatment with copper and zinc, *R. toruloides* increases expression of anaplerosis via fatty acid and branched-chain amino acid catabolism concomitant with increases in expression of cysteine and glutathione biosynthesis pathways.** Metabolic pathway map displaying the putative pathways in *R. toruloides* for branched-chain amino acid catabolism, fatty acid catabolism via acylcarnitine, the citric acid cycle, the glyoxylate cycle, the methylcitrate cycle, sulfate assimilation, and cysteine/glutathione biosynthesis. Each reaction is labeled with solid arrows indicating directionality, the name of the enzyme catalyzing the reaction, the *R. toruloides* gene ID (excluding the "RTO4_" prefix) encoding the enzyme, and, in parentheses, the $\log_2$(fold change) in expression for the gene in 5 mM copper sulfate plus 4.78 mM zinc sulfate relative to ammonium-phosphate buffer. Gene IDs, $\log_2$(fold change), and arrows are: bolded and colored red if the gene was at least two-fold increased in expression ($p_{adj} < 0.05$); colored black if the gene was not significantly differentially expressed by at least two-fold; or bolded and colored blue if the gene was at least two-fold decreased in expression ($p_{adj} < 0.05$). Dashed arrows indicate connections of metabolites within and between pathways. A "?" is annotated next to 11331 in the methylcitrate synthase reaction because it is unclear if *R. toruloides* catalyzes both citrate and methylcitrate synthesis via the same enzyme or if *R. toruloides* has a dedicated methylcitrate synthase that is unannotated. The following abbreviations are used for enzymes and metabolites: BCAT is branched-chain aminotransferase, BCKD is branched-chain keto acid dehydrogenase, ACAT is Acetyl-CoA acetyltransferase, MCCC is Methylcrotonyl-CoA carboxylase, HSD17β10 is 17-β-hydroxysteroid dehydrogenase 10, HIBCH is 3-hydroxyisobutyryl-CoA hydrolase, HIBDH is 3-hydroxyisobutyrate dehydrogenase, OAH/OAS is O-acetyl homoserine-O-acetyl serine sulfhydrylase, HMG is 3-hydroxy-3-methylglutaryl, APS is adenosine 5'-phosphosulfate, PAPS is 3'-phosphoadenylyl-sulfate.

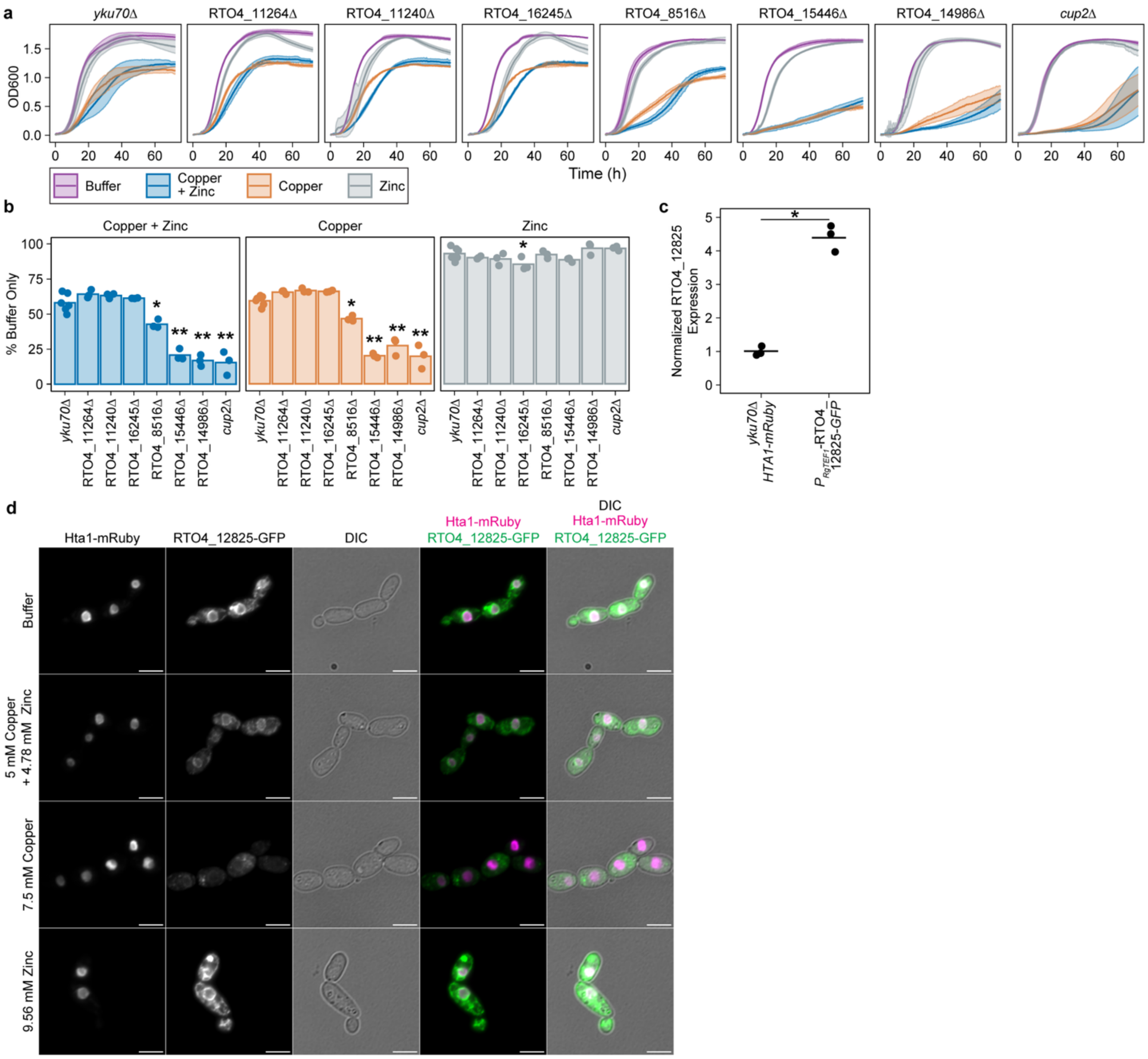


**Fig S9. *R. toruloides* genes differentially expressed in response to copper and zinc are involved in copper and zinc resistance. (A)** Growth (OD600) of the *R. toruloides yku70*Δ parental strain and indicated deletion mutant strains grown on media supplemented with ammonium-phosphate buffer or ammonium-phosphate buffer with 5 mM copper sulfate plus 4.78 mM zinc sulfate, 7.5 mM copper sulfate, or 9.56 mM zinc sulfate. Lines are averages and colored bands are standard deviations of at least three biological replicates. **(B)** Area under the curve from (**A**) of the indicated cells grown in the indicated conditions normalized to the area under the curve of the same strain on media supplemented with ammonium-phosphate buffer.

Bars are averages and dots are individual data points of at least three biological replicates. **(C)** Reverse transcription quantitative PCR (RT-qPCR) analysis of RTO4_12825 expression in the *yku70*Δ *HTA1-mRuby2* parental strain and $P_{RgTEF1}$-RTO4_12825-*GFP* cells grown in minimal media. Expression is normalized to the average expression of RTO4_12825 in the *yku70*Δ *HTA1-mRuby2* parental strain. **(D)** $P_{RgTEF1}$-RTO4_12825-*GFP* cells were exposed media supplemented with ammonium-phosphate buffer or ammonium-phosphate buffer with 5 mM copper sulfate plus 4.78 mM zinc sulfate, 7.5 mM copper sulfate, or 9.56 mM zinc sulfate for eight hours. Cells were fixed prior to imaging. Fluorescent microscopy was performed using a Nikon Eclipse Ti2-E inverted microscope equipped with a prime BSI express CMOS camera using a CFI Plan Apochromat 100x oil objective. The scale bar is 5 µm. Images are representative of three biological replicates. Lines are averages and dots are individual data points of three biological replicates. *$p < 0.05$ and **$p < 10^{-8}$ as determined by (**B**) a one-way ANOVA with a Dunnett's post-hoc test compared to the *yku70*Δ parental strain or (**C**) a Welch's t-test.

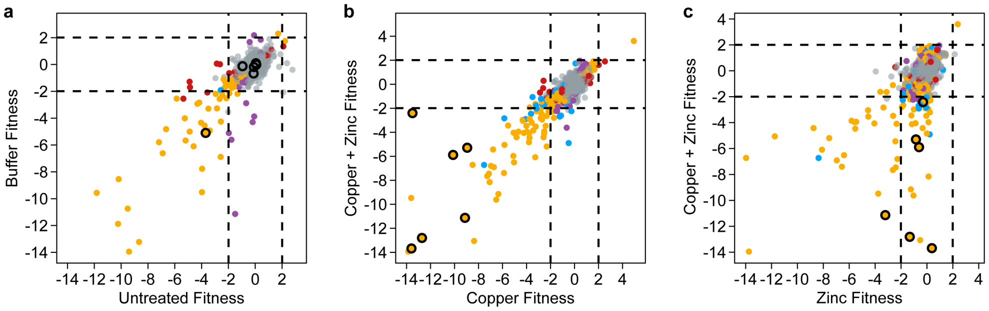


| | | | | | |
|---|---|---|---|---|---|
| x-axis condition vs y-axis condition | **$p_{adj}$ < 0.05** | $p_{adj} \geq 0.05$ | $p_{adj} \geq 0.05$ | $p_{adj} \geq 0.05$ | $p_{adj} \geq 0.05$ |
| x-axis condition vs starting population | NA | **$p_{adj}$ < 0.05** | **$p_{adj}$ < 0.05** | $p_{adj} \geq 0.05$ | $p_{adj} \geq 0.05$ |
| y-axis condition vs starting population | NA | **$p_{adj}$ < 0.05** | $p_{adj} \geq 0.05$ | **$p_{adj}$ < 0.05** | $p_{adj} \geq 0.05$ |
| Confirmed phenotype: O | | | | | |

**Fig S10. *R. toruloides* genes have similar fitness phenotypes during exposure to copper alone and copper in combination with zinc. (A)** Gene fitness scores obtained by growing a barcoded mutant library[5] on media (untreated) or media supplemented with ammonium-phosphate buffer. **(B-C)** Gene fitness scores obtained by growing a barcoded mutant library[5] on media with ammonium-phosphate buffer and 3.33 mM copper sulfate plus 3.19 mM zinc sulfate versus (**B**) 6.29 mM copper sulfate or (**C**) 9 mM zinc sulfate. Genes with significant fitness scores ($p_{adj} < 0.05$) in the x-axis condition relative to the y-axis condition are indicated in blue; in both the x-axis and y-axis conditions relative to the starting population are indicated in yellow; in only the x-axis condition relative to the starting population are indicated in red; in only the y-axis condition relative to the starting population are indicated in purple. Genes without a significant fitness score in the x-axis or y-axis condition relative to the starting population or the x-axis condition relative to the y-axis condition ($p_{adj} \geq 0.05$) are indicated in grey. Genes with phenotypes confirmed by a clean deletion strain are circled in black.

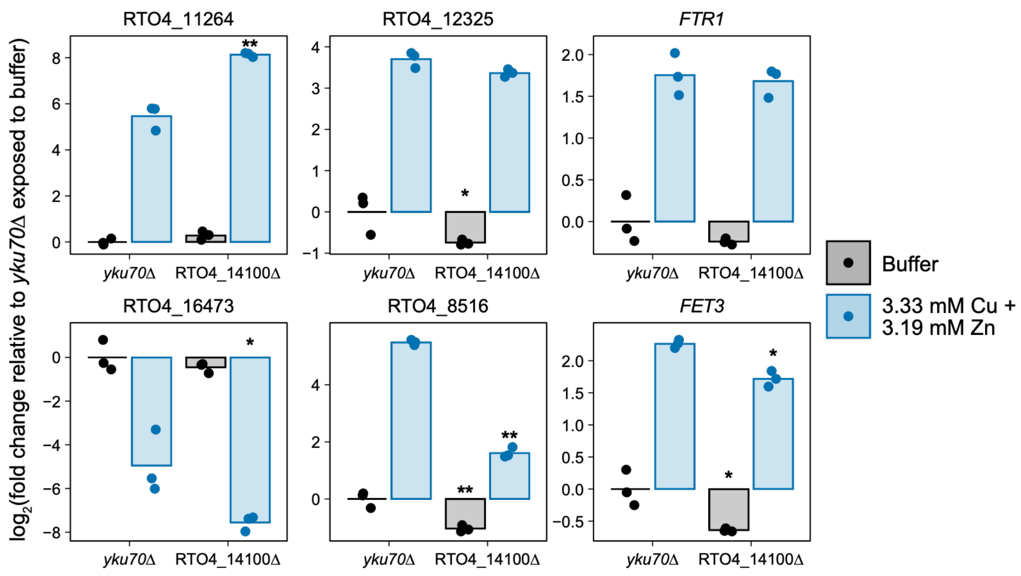


**Fig S11. RTO4_14100 regulates a subset of genes in the *R. toruloides* transcriptional response to copper and zinc.** Expression of the indicated genes in *yku70*Δ (parental strain) and RTO4_14100Δ cells in media supplemented with ammonium-phosphate buffer or ammonium-phosphate buffer with 3.33 mM copper sulfate plus 3.19 mM zinc sulfate as determined by RT-qPCR. Gene expression was normalized to the expression of the indicated gene in the *yku70*Δ parental strain treated with ammonium-phosphate buffer. *p < 0.05 and **p < 0.001 for RTO4_14100Δ cells compared to *yku70*Δ cells in the same condition, as determined by two-way ANOVA with a Tukey's post-hoc test.

### SI TABLES

**Table S1. *Neurospora crassa* strains used in this study.**

| Strain name | Genotype | Source |
|---|---|---|
| Wild type | Wild type *mat A* | FGSC* 2489[4] |
| Wild type | Wild type *mat a* | FGSC 4200[4] |
| ΔNCU00144 *mat A* | ΔNCU00144::*Hyg*$^{R}$** *mat A* | FGSC 11339[3] |
| ΔNCU00144 *mat a* | ΔNCU00144::*Hyg*$^{R}$ *mat a* | FGSC 11340[3] |
| ΔNCU00227 *mat A* | ΔNCU00227::*Hyg*$^{R}$ *mat A* | FGSC 15857[3] |
| ΔNCU00227 *mat a* | ΔNCU00227::*Hyg*$^{R}$ *mat a* | FGSC 15856[3] |
| ΔNCU01138 *mat A* | ΔNCU01138::*Hyg*$^{R}$ *mat A* | FGSC 13030[3] |
| ΔNCU01138 *mat a* | ΔNCU01138::*Hyg*$^{R}$ *mat a* | FGSC 13029[3] |
| Δ*uce-12 mat A* | Δ*uce-12* (NCU01188)::*Hyg*$^{R}$ *mat A* | FGSC 16552[3] |
| Δ*gst-3 mat a* | Δ*gst-3* (NCU01320)::*Hyg*$^{R}$ *mat a* | FGSC 11528[3] |
| Δ*aga-2 mat a* | Δ*aga-2* (NCU02084)::*Hyg*$^{R}$ *mat a* | FGSC 16048[3] |
| Δ*cfp mat a* | Δ*cfp* (NCU02193)::*Hyg*$^{R}$ *mat a* | FGSC 18065[3] |
| Δ*cfp mat A* | Δ*cfp*::*Hyg*$^{R}$ *mat A* | FGSC 18066[3] |
| Δ*mig-8 mat A* | Δ*mig-8* (NCU03369)::*Hyg*$^{R}$ *mat A* | FGSC 14512[3] |
| Δ*trx-5 mat a* | Δ*trx-5* (NCU03714)::*Hyg*$^{R}$ *mat a* | FGSC 12993[3] |
| Δ*trx-5 mat A* | Δ*trx-5*::*Hyg*$^{R}$ *mat A* | FGSC 12994[3] |
| Δ*znf-27 mat A* | Δ*znf-27* (NCU04022)::*Hyg*$^{R}$ *mat A* | FGSC 17880[3] |
| Δ*trm-47 mat a* | Δ*trm-47* (NCU04076)::*Hyg*$^{R}$ *mat a* | FGSC 11533[3] |
| Δ*hne-1 mat A* | Δ*hne-1* (NCU04172)::*Hyg*$^{R}$ *mat A* | FGSC 11662[3] |
| ΔNCU04645 *mat a* | ΔNCU04645::*Hyg*$^{R}$ *mat a* | FGSC 19133[3] |
| Δ*aap-7 mat a* | Δ*aap-7* (NCU04942)::*Hyg*$^{R}$ *mat a* | FGSC 13329[3] |
| Δ*aap-7 mat A* | Δ*aap-7*::*Hyg*$^{R}$ *mat A* | FGSC 13330[3] |
| Δ*gst-1 mat a* | Δ*gst-1* (NCU05780)::*Hyg*$^{R}$ *mat a* | FGSC 14007[3] |
| Δ*gst-1* mat A | Δ*gst-1*::*Hyg*$^{R}$ *mat A* | FGSC 14008[3] |
| Δ*adh-5 mat A* | Δ*adh-5* (NCU06303)::*Hyg*$^{R}$ *mat A* | FGSC 19999[3] |
| Δ*adh-5 mat A* | Δ*adh-5* (NCU06303)::*Hyg*$^{R}$ *mat A* | FGSC 20000[3] |
| Δ*znf-2 mat a* | Δ*znf-2* (NCU06586)::*Hyg*$^{R}$ *mat a* | FGSC 20201[3] |
| Δ*znf-2 mat A* | Δ*znf-2*::*Hyg*$^{R}$ *mat A* | FGSC 20202[3] |
| Δ*hsp30-2 mat a* | Δ*hsp30-2* (NCU07232)::*Hyg*$^{R}$ *mat a* | FGSC 11568[3] |
| Δ*mig-2 mat a* | Δ*mig-2* (NCU07452)::*Hyg*$^{R}$ *mat a* | FGSC 14184[3] |
| Δ*abc-1 mat A* | Δ*abc-1* (NCU08056)::*Hyg*$^{R}$ *mat A* | FGSC 13048[3] |
| Δ*adh-8 mat A* | Δ*adh-8* (NCU08402)::*Hyg*$^{R}$ *mat A* | FGSC 23476[3] |
| Δ*rbg-43 mat a* | Δ*rbg-43* (NCU08663)::*Hyg*$^{R}$ *mat a* | FGSC 15507[3] |
| ΔNCU08900 *mat a* | ΔNCU08900::*Hyg*$^{R}$ *mat a* | FGSC 18417[3] |
| ΔNCU08900 *mat A* | ΔNCU08900::*Hyg*$^{R}$ *mat A* | FGSC 18418[3] |
| Δ*cys-16 mat A* | Δ*cys-16* (NCU09230)::*Hyg*$^{R}$ *mat A* | FGSC 22142[3] |
| Δ*hsp30 mat a* | Δ*hsp30* (NCU09364)::*Hyg*$^{R}$ *mat a* | FGSC 11267[3] |
| Δ*mig-12 mat a* | Δ*mig-12* (NCU09830)::*Hyg*$^{R}$ *mat a* | FGSC 11262[3] |
| Δ*mig-12 mat A* | Δ*mig-12*::*Hyg*$^{R}$ *mat A* | FGSC 11263[3] |
| Δ*gst-4 mat a* | Δ*gst-4* (NCU10521)::*Hyg*$^{R}$ *mat a* | FGSC 18656[3] |
| Δ*gst-4 mat A* | Δ*gst-4*::*Hyg*$^{R}$ *mat A* | FGSC 18657[3] |

*FGSC stands for Fungal Genetics Stock Center[4]

***Hyg*$^{R}$ stands for hygromycin resistance cassette

**Table S2. *Rhodotorula (Rhodosporidium) toruloides* strains used in this study.**

| Strain name | Genotype | Source |
|---|---|---|
| Wild type | Wild type IFO0880 (also known as NBRC 0880) | NBRC* culture collection |
| *yku70*Δ | *yku70*(RTO4_11920)Δ::$Hyg^R$** | [6] |
| RTO4_8516Δ | RTO4_8516Δ::$Nat^R$*** *yku70*Δ::$Hyg^R$ | This study |
| RTO4_8997Δ | RTO4_8997Δ::$Nat^R$ *yku70*Δ::$Hyg^R$ | This study |
| *vps22*Δ | *vps22* (RTO4_9066)Δ::$Nat^R$ *yku70*Δ::$Hyg^R$ | This study |
| *fet3*Δ | *fet3* (RTO4_9364)Δ::$Nat^R$ *yku70*Δ::$Hyg^R$ | This study |
| RTO4_10354Δ | RTO4_10354Δ::$Nat^R$ *yku70*Δ::$Hyg^R$ | This study |
| *vps35*Δ | *vps35* (RTO4_10531)Δ::$Nat^R$ *yku70*Δ::$Hyg^R$ | This study |
| *pcs2*Δ | *pcs2* (RTO4_10995)Δ::$Nat^R$ *yku70*Δ::$Hyg^R$ | This study |
| RTO4_11240Δ | RTO4_11240Δ::$Nat^R$ *yku70*Δ::$Hyg^R$ | This study |
| RTO4_11264Δ | RTO4_11264Δ::$Nat^R$ *yku70*Δ::$Hyg^R$ | This study |
| *mcy1*Δ | *mcy1* (RTO4_12031)Δ::$Nat^R$ *yku70*Δ::$Hyg^R$ | This study |
| RTO4_12825Δ | RTO4_12825Δ::$Nat^R$ *yku70*Δ::$Hyg^R$ | This study |
| *cup2*Δ | *cup2* (RTO4_14231)Δ::$Nat^R$ *yku70*Δ::$Hyg^R$ | This study |
| RTO4_14100 | RTO4_14100Δ::$Nat^R$ *yku70*Δ::$Hyg^R$ | This study |
| RTO4_14986Δ | RTO4_14986Δ::$Nat^R$ *yku70*Δ::$Hyg^R$ | This study |
| RTO4_15446Δ | RTO4_15446Δ::$Nat^R$ *yku70*Δ::$Hyg^R$ | This study |
| *ftr1*Δ | *ftr1* (RTO4_15844)Δ::$Nat^R$ *yku70*Δ::$Hyg^R$ | This study |
| RTO4_16245Δ | RTO4_16245Δ::$Nat^R$ *yku70*Δ::$Hyg^R$ | This study |
| *HTA1-mRuby2* | *HTA1*(RTO4_12090)*-mRuby2 yku70*Δ::$Hyg^R$ | This study |
| *HTA1-mRuby2*<br>RTO4_12825-*mEGFP* | *HTA1-mRuby2*<br>*yku70*Δ::$P_{RgTEF1}$-RTO4_12825-*mEGFP*:$G418^R$**** | This study |

*NBRC stands for National Institute of Technology and Evaluation (NITE) Biological Resource Center

**$Hyg^R$ stands for hygromycin resistance cassette

***$Nat^R$ stands for nourseothricin resistance cassette

****$G418^R$ stands for G418 resistance cassette

**Table S3. Primers used in this study.**

| Primer use | Forward primer 5' → 3' | Reverse primer 5' → 3' |
|---|---|---|
| BarSeq T0 Replicate 1 | CAAGCAGAAGACGGCATACGAGATCGTGATGTGACTGGAGTTCAGACGTGTGCTCTTCCGATCTGATGTCCACGAGGTCTCT | AATGATACGGCGACCACCGAGATCTACACTCTTTCCCTACACGACGCTCTTCCGATCTNGCACTAGTCGACCTGCAGCGTACG |
| BarSeq T0 Replicate 2 | CAAGCAGAAGACGGCATACGAGATACATCGGTGACTGGAGTTCAGACGTGTGCTCTTCCGATCTGATGTCCACGAGGTCTCT | AATGATACGGCGACCACCGAGATCTACACTCTTTCCCTACACGACGCTCTTCCGATCTNNTGTAGCGTCGACCTGCAGCGTACG |
| BarSeq T0 Replicate 3 | CAAGCAGAAGACGGCATACGAGATGCCTAAGTGACTGGAGTTCAGACGTGTGCTCTTCCGATCTGATGTCCACGAGGTCTCT | AATGATACGGCGACCACCGAGATCTACACTCTTTCCCTACACGACGCTCTTCCGATCTNNNCGGATTGTCGACCTGCAGCGTACG |
| BarSeq no treatment Replicate 1 | CAAGCAGAAGACGGCATACGAGATTGGTCAGTGACTGGAGTTCAGACGTGTGCTCTTCCGATCTGATGTCCACGAGGTCTCT | AATGATACGGCGACCACCGAGATCTACACTCTTTCCCTACACGACGCTCTTCCGATCTNNNNACCAGTGTCGACCTGCAGCGTACG |
| BarSeq no treatment Replicate 2 | CAAGCAGAAGACGGCATACGAGATCACTGTGTGACTGGAGTTCAGACGTGTGCTCTTCCGATCTGATGTCCACGAGGTCTCT | AATGATACGGCGACCACCGAGATCTACACTCTTTCCCTACACGACGCTCTTCCGATCTNGTGACAGTCGACCTGCAGCGTACG |
| BarSeq no treatment Replicate 3 | CAAGCAGAAGACGGCATACGAGATATTGGCGTGACTGGAGTTCAGACGTGTGCTCTTCCGATCTGATGTCCACGAGGTCTCT | AATGATACGGCGACCACCGAGATCTACACTCTTTCCCTACACGACGCTCTTCCGATCTNNTAACCGGTCGACCTGCAGCGTACG |
| BarSeq ammonium-phosphate buffer Replicate 1 | CAAGCAGAAGACGGCATACGAGATGATCTGGTGACTGGAGTTCAGACGTGTGCTCTTCCGATCTGATGTCCACGAGGTCTCT | AATGATACGGCGACCACCGAGATCTACACTCTTTCCCTACACGACGCTCTTCCGATCTNNNCTAGACGTCGACCTGCAGCGTACG |
| BarSeq ammonium-phosphate buffer Replicate 2 | CAAGCAGAAGACGGCATACGAGATTCAAGTGTGACTGGAGTTCAGACGTGTGCTCTTCCGATCTGATGTCCACGAGGTCTCT | AATGATACGGCGACCACCGAGATCTACACTCTTTCCCTACACGACGCTCTTCCGATCTNNNNAGTTCAGTCGACCTGCAGCGTACG |
| BarSeq ammonium-phosphate buffer Replicate 3 | CAAGCAGAAGACGGCATACGAGATCTGATCGTGACTGGAGTTCAGACGTGTGCTCTTCCGATCTGATGTCCACGAGGTCTCT | AATGATACGGCGACCACCGAGATCTACACTCTTTCCCTACACGACGCTCTTCCGATCTNGACTAGGTCGACCTGCAGCGTACG |
| BarSeq 3.33 mM copper sulfate plus 3.19 mM zinc sulfate Replicate 1 | CAAGCAGAAGACGGCATACGAGATAAGCTAGTGACTGGAGTTCAGACGTGTGCTCTTCCGATCTGATGTCCACGAGGTCTCT | AATGATACGGCGACCACCGAGATCTACACTCTTTCCCTACACGACGCTCTTCCGATCTNNTTCGATGTCGACCTGCAGCGTACG |
| BarSeq 3.33 mM copper sulfate plus 3.19 mM zinc sulfate Replicate 2 | CAAGCAGAAGACGGCATACGAGATGTAGCCGTGACTGGAGTTCAGACGTGTGCTCTTCCGATCTGATGTCCACGAGGTCTCT | AATGATACGGCGACCACCGAGATCTACACTCTTTCCCTACACGACGCTCTTCCGATCTNNNCATCGGGTCGACCTGCAGCGTACG |
| BarSeq 3.33 mM copper sulfate | CAAGCAGAAGACGGCATACGAGATTACAAGGTGACTGGAGTTCAGA | AATGATACGGCGACCACCGAGATCTACACTCTTTCCCTACACG |

| | | |
|---|---|---|
| plus 3.19 mM zinc sulfate Replicate 3 | CGTGTGCTCTTCCGATCTGATGTCCACGAGGTCTCT | ACGCTCTTCCGATCTNNNNATGTTCGTCGACCTGCAGCGTACG |
| BarSeq 6.29 mM copper sulfate Replicate 1 | CAAGCAGAAGACGGCATACGAGATTTGACTGTGACTGGAGTTCAGACGTGTGCTCTTCCGATCTGATGTCCACGAGGTCTCT | AATGATACGGCGACCACCGAGATCTACACTCTTTCCCTACACGACGCTCTTCCGATCTNAACTGAGTCGACCTGCAGCGTACG |
| BarSeq 6.29 mM copper sulfate Replicate 2 | CAAGCAGAAGACGGCATACGAGATGGAACTGTGACTGGAGTTCAGACGTGTGCTCTTCCGATCTGATGTCCACGAGGTCTCT | AATGATACGGCGACCACCGAGATCTACACTCTTTCCCTACACGACGCTCTTCCGATCTNNCCTTGAGTCGACCTGCAGCGTACG |
| BarSeq 6.29 mM copper sulfate Replicate 3 | CAAGCAGAAGACGGCATACGAGATTGACATGTGACTGGAGTTCAGACGTGTGCTCTTCCGATCTGATGTCCACGAGGTCTCT | AATGATACGGCGACCACCGAGATCTACACTCTTTCCCTACACGACGCTCTTCCGATCTNNNACTGTAGTCGACCTGCAGCGTACG |
| BarSeq 9 mM zinc sulfate Replicate 1 | CAAGCAGAAGACGGCATACGAGATGGACGGGTGACTGGAGTTCAGACGTGTGCTCTTCCGATCTGATGTCCACGAGGTCTCT | AATGATACGGCGACCACCGAGATCTACACTCTTTCCCTACACGACGCTCTTCCGATCTNNNNCCTGCCGTCGACCTGCAGCGTACG |
| BarSeq 9 mM zinc sulfate Replicate 2 | CAAGCAGAAGACGGCATACGAGATCTCTACGTGACTGGAGTTCAGACGTGTGCTCTTCCGATCTGATGTCCACGAGGTCTCT | AATGATACGGCGACCACCGAGATCTACACTCTTTCCCTACACGACGCTCTTCCGATCTNGAGATGGTCGACCTGCAGCGTACG |
| BarSeq 9 mM zinc sulfate Replicate 3 | CAAGCAGAAGACGGCATACGAGATGCGGACGTGACTGGAGTTCAGACGTGTGCTCTTCCGATCTGATGTCCACGAGGTCTCT | AATGATACGGCGACCACCGAGATCTACACTCTTTCCCTACACGACGCTCTTCCGATCTNNCGCCTGGTCGACCTGCAGCGTACG |
| RTO4_12825 RT-qPCR | CTTTGTACGCCGATTGGGATGG | GTAGAGGAGGATGCCAGACGA |
| *FET3* RT-qPCR | GGATCTTCCATTGCCACATTCA | ATCCGTGACAGACATTTTGCC |
| *FTR1* RT-qPCR | ATCAGGGAGAAGAAGGAGATTG | TAAGTCCACCATAACACCCTTA |
| RTO4_12325 RT-qPCR | ATCCTACTTCTACACCTACCTC | CGGACGACGATGCCCATTAC |
| RTO4_11264 RT-qPCR | GTCTTCACGAAGGACAAGCAAGAG | GCATCGAAGTCGGCATGAACC |
| RTO4_16473 RT-qPCR | CTCTCTCGCCCACAGGAGAAG | CGGAGTAGATGGCGTAGGTGG |
| RTO4_8516 RT-qPCR | GCGGCTTTGCGAATGCCTC | CAGCAGATGAAGGTGCGGTTC |
| RTO4_9189 RT-qPCR | TCGAGGGCGAATGGCTCAAG | GCACGACAAGCCTTCGGATC |
| *ACT1* RT-qPCR | CGTCCTCTCGCTCTATGCCTC | CTTGATGAGGTAGTCGGTCAGGTC |

**SI DATASETS**

**Dataset S1. Expression of all genes following an eight hour exposure of *N. crassa* mycelia to media, media supplemented with ammonium-phosphate buffer alone, or media supplemented with ammonium-phosphate buffer and 3.33 mM copper sulfate plus 3.19 mM zinc sulfate, 2 mM copper sulfate plus 1.91 mM zinc sulfate, 5 mM copper sulfate, 3.33 mM copper sulfate, 3.19 mM zinc sulfate, or 4.78 mM zinc sulfate.**

**Dataset S2. Differential gene expression calculated by DESeq2[2] in the indicated comparisons for *N. crassa* mycelia exposed to media, media supplemented with ammonium-phosphate buffer alone, or media supplemented with ammonium-phosphate buffer and 3.33 mM copper sulfate plus 3.19 mM zinc sulfate, 2 mM copper sulfate plus 1.91 mM zinc sulfate, 5 mM copper sulfate, 3.33 mM copper sulfate, 3.19 mM zinc sulfate, or 4.78 mM zinc sulfate for eight hours.**

**Dataset S3. Expression of all genes following an eight hour exposure of *R. toruloides* cells to media, media supplemented with ammonium-phosphate buffer alone, or media supplemented with ammonium-phosphate buffer and 5 mM copper sulfate plus 4.78 mM zinc sulfate, 3.33 mM copper sulfate plus 3.19 mM zinc sulfate, 6.29 mM copper sulfate, 3.33 mM copper sulfate, or 9 mM zinc sulfate.**

**Dataset S4. Differential gene expression calculated by DESeq2[2] in the indicated comparisons for *R. toruloides* cells exposed to media, media supplemented with ammonium-phosphate buffer alone, or media supplemented with ammonium-phosphate buffer and 5 mM copper sulfate plus 4.78 mM zinc sulfate, 3.33 mM copper sulfate plus 3.19 mM zinc sulfate, 6.29 mM copper sulfate, 3.33 mM copper sulfate, or 9 mM zinc sulfate for eight hours.**

**Dataset S5. Gene fitness scores and $p_{adj}$-values of the *R. toruloides* barcoded mutant library grown in media, media supplemented with ammonium-phosphate buffer alone, or media supplemented with ammonium-phosphate buffer and 3.33 mM copper sulfate plus 3.19 mM zinc sulfate, 6.3 mM copper sulfate, or 9 mM zinc.**

**Dataset S6. Numerical values used to generate all bar and line graphs.**

### SI REFERENCES


1. Zheng, S. *et al.* SynergyFinder Plus: Toward Better Interpretation and Annotation of Drug Combination Screening Datasets. *Genomics Proteomics Bioinformatics* **20**, 587–596 (2022).
2. Love, M. I., Huber, W. & Anders, S. Moderated estimation of fold change and dispersion for RNA-seq data with DESeq2. *Genome Biol.* **15**, 550 (2014).
3. Colot, H. V. *et al.* A high-throughput gene knockout procedure for Neurospora reveals functions for multiple transcription factors. *Proc. Natl. Acad. Sci.* **103**, 10352–10357 (2006).
4. McCluskey, K., Wiest, A. & Plamann, M. The Fungal Genetics Stock Center: a repository for 50 years of fungal genetics research. *J. Biosci.* **35**, 119–126 (2010).
5. Coradetti, S. T. *et al.* Functional genomics of lipid metabolism in the oleaginous yeast Rhodosporidium toruloides. *eLife* **7**, e32110 (2018).
6. Zhang, S., Ito, M., Skerker, J. M., Arkin, A. P. & Rao, C. V. Metabolic engineering of the oleaginous yeast Rhodosporidium toruloides IFO0880 for lipid overproduction during high-density fermentation. *Appl. Microbiol. Biotechnol.* **100**, 9393–9405 (2016).
